# High-resolution single cell transcriptome analysis of zebrafish sensory hair cell regeneration

**DOI:** 10.1101/2021.07.15.452338

**Authors:** Sungmin Baek, Nhung T. T. Tran, Daniel C. Diaz, Ya-Yin Tsai, Joaquin Navajas Acedo, Mark E. Lush, Tatjana Piotrowski

## Abstract

Loss of sensory hair cells in the mammalian inner ear leads to permanent hearing and vestibular defects, whereas loss of hair cells in zebrafish results in their regeneration. We used scRNA-Seq to characterize the transcriptional dynamics of hair cell regeneration in zebrafish at unprecedented spatio-temporal resolution. We uncovered three, sequentially activated modules. First, an injury/inflammatory response and downregulation of progenitor/stem cell maintenance genes within minutes after hair cell loss. Second, the transient activation of regeneration-specific genes. And third, a robust reactivation of developmental gene programs, including hair cell specification, cell cycle activation, ribosome biogenesis, and a metabolic switch to oxidative phosphorylation. The results are not only relevant for our understanding of hair cell regeneration and how we might be able to trigger it in mammals but also for regenerative processes in general. The data is searchable and publicly accessible via a web-based interface.

## Introduction

Humans in particular, and mammals in general, fail to regenerate mechanosensory hair cells in their inner ears. Noise, drug or age-induced hearing loss due to hair cell death is, therefore, permanent (Furness, 2015; Wagner and Shin, 2019; Wong and Ryan, 2015). In contrast, fish, chicken and amphibians constantly replace dying hair cells through proliferation and differentiation of support cells (Corwin and Cotanche, 1988; Corwin and Warchol, 1991; Cruz et al., 2015; Pinto-Teixeira et al., 2015; Ryals and Rubel, 1988). Because neonatal mouse support cells are able to produce a modest number of hair cells after injury and some vestibular hair cells turn over spontaneously, there is hope that elucidating the gene regulatory networks underpinning hair cell regeneration, may lead to the restoration of hearing loss. (Bucks et al., 2017; Burns et al., 2012; Burns and Stone, 2017; Cox et al., 2014; Du et al., 2013; Forge et al., 1993; Korrapati et al., 2013; Warchol and Corwin, 1996; White et al., 2006). One strategy to identify candidate genes and pathways that might trigger hair cell regeneration in mammals has been to identify genes involved in hair cell development (Burns et al., 2015; Cai and Groves, 2015; Jen et al., 2019; Lee et al., 2020; Sayyid et al., 2019; Scheffer et al., 2015; Valdivia et al., 2011; Zhu et al., 2019). Another promising strategy is to identify the molecular underpinnings of hair cell regeneration in regenerating species, such as chicken and zebrafish and use this blueprint of regeneration to experimentally trigger regeneration in mammals (Behra et al., 2009; Jiang et al., 2014; Ku et al., 2014; Pei et al., 2018; Roccio et al., 2018; Scheibinger et al., 2018; Steiner et al., 2014). However, the number of regenerated hair cells and hearing improvement attained in mammals thus far has been modest. This is likely due to the limited number of known candidate genes and our general lack of understanding of hair cell type specific expression dynamics at a fine time scale.

The zebrafish sensory lateral line is a powerful model to study hair cell development and regeneration because of its accessibility and relative ease of experimental manipulation (Figure 1A; (Denans et al., 2019; Kniss et al., 2016; Lush and Piotrowski, 2014; Ma and Raible, 2009; Romero-Carvajal et al., 2015; Viader-Llargues et al., 2018)). Zebrafish and mammalian sensory hair cells are very similar, and loss of lateral line hair cell genes does not only cause loss of hair cell function in fish, but also causes deafness and vestibular dysfunction in mammals (Nicolson, 2005; Whitfield, 2002).

**Figure 1.**
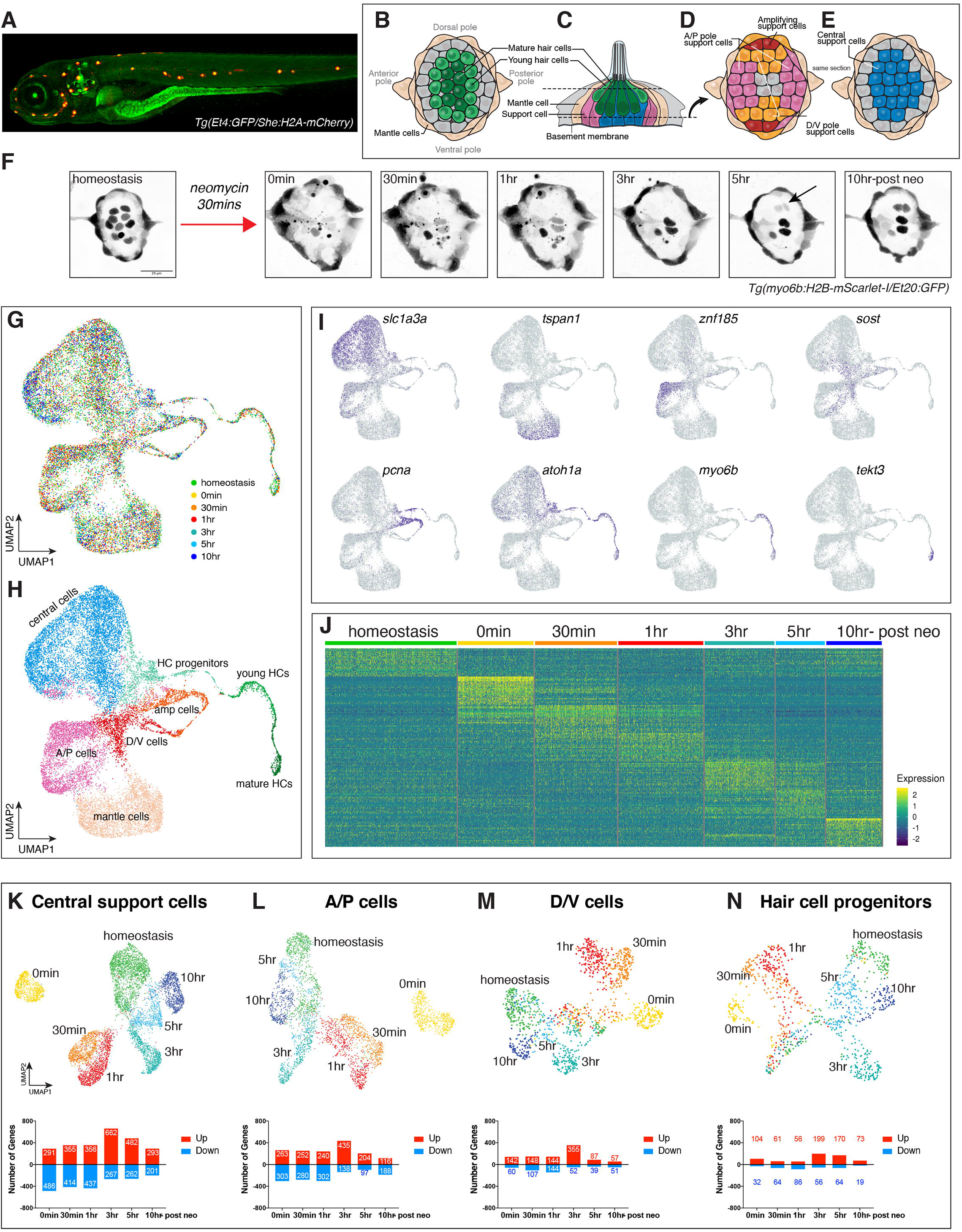
Spatio-temporal scRNA-Seq analysis during lateral line hair cell regeneration. (A) 5dpf *Tg(she:H2A-mCherry)* larvae were used for FACS of lateral line cells. (B - E) Illustration of homeostatic lateral line cell types. (B, D, E) dorsal view and (C) lateral view. (F) Confocal images of *Tg(myo6b:H2B-mScarlet-I/et20:GFP)* neuromasts during hair cell regeneration. Scale bar: 20μm, arrow indicates two new hair cells. (G) Integration of scRNA-Seq samples of seven different time points in UMAP latent space. (H) UMAP of a cluster analysis identifies eight different neuromast cell types. (I) Feature plots of cell type-specific marker genes. (J) Heatmap visualizing the top 25 genes in each timepoint during hair cell regeneration. (Differential gene expression analysis of each time point vs. all other time points) (K - N) UMAPs of subsetted (K) central support cells, (L) A/P cells, (M) D/V cells and (N) hair cell progenitors with bar graphs showing the number of up and down-regulated genes during regeneration (differential gene expression analysis of each time point vs. homeostasis in individual cell type).

We have previously performed bulk RNA-Seq analyses of regenerating lateral line hair cells that identified candidate genes and pathways. However, this study lacked spatio-temporal resolution (Jiang et al., 2014). To identify the cell populations involved in regeneration, we analyzed the different cell populations that respond to hair cell death in time lapse and BrdU analyses and have characterized all cell populations in homeostatic lateral line sensory organs (neuromasts) using scRNA-Seq (Figures 1B-E; (Lush et al., 2019; Romero-Carvajal et al., 2015)). Lateral line sensory organs consist of sensory hair cells surrounded by support cells and an outermost ring of mantle cells. Support cells can be subdivided into anterior-posterior (A/P), dorso-ventral (D/V) and a central sub population (Romero-Carvajal et al., 2015; Thomas and Raible, 2019; Wibowo et al., 2011). During regeneration support cells divide before differentiating into hair cells (López-Schier and Hudspeth, 2006; Ma et al., 2008; Mackenzie and Raible, 2012; Wibowo et al., 2011). Central support cells are the immediate precursors of hair cells (Lush et al., 2019; Romero-Carvajal et al., 2015). D/V and A/P support cells also give rise to hair cells but it is not known if they first acquire a central support cell fate or if they directly differentiate into hair cells during regeneration. Mantle cells do not respond behaviorally to hair cell loss, but they generate support cells and hair cells after severe injury to the sensory organ (Denans et al., 2019; Romero-Carvajal et al., 2015; Thomas and Raible, 2019; Viader-Llargues et al., 2018).

Previous studies have shown that all regenerative cell divisions are symmetric. Central and A/P support cells divide and give rise to two hair cells, whereas D/V cells that are in contact with mantle cells (amplifying cells) also divide symmetrically but do not differentiate and instead give rise to two support cells (Romero-Carvajal et al., 2015). Manipulations of the Notch and Wnt signaling pathways demonstrated that the majority, or maybe even all support cell populations can be induced to proliferate and either give rise to only hair cells or to only support cells. We therefore concluded that the majority of support cells are competent to serve as hair cell progenitors.

Faithful regeneration of sensory cells requires the precise orchestration of stem/progenitor cell activation, proliferation, cell lineage determination and differentiation. Of all the vertebrate regeneration models studied, such as fin, heart and retina regeneration, hair cell regeneration is by far the fastest. Zebrafish lateral line cells die within 30min of killing hair cells and the first newly regenerated hair cells arise around 5 hours later. Differential gene expression, enrichment term analysis, pseudotime analysis, and validation by in situ hybridization allowed us to define the molecular changes taking place during hair cell regeneration at unprecedented levels of resolution. We correlated the transcriptional dynamics with previously defined cellular behaviors to build the foundation for the establishment of a gene regulatory network (GRN). We discovered three, sequentially activated modules. The first module consists of a systemic injury/inflammatory response, which likely plays a crucial role in the initiation of regeneration (Beisaw et al., 2020; Wang et al., 2020). The second module is characterized by a transient activation of regeneration-specific genes. The longest, third phase of regeneration recapitulates developmental events. Thus, once initiated, regeneration recapitulates development but the early injury/inflammatory response is regeneration-specific. The resulting gene expression dynamics are publicly available and readily searchable via web-based interfaces: https://piotrowskilab.shinyapps.io/neuromast_regeneration_scRNA-Seq_pub_2021/ and gEAR (https://umgear.org/p?l=e0d00950) (Orvis et al., 2021).

Knowing when and in which cell types genes and pathways are modulated, and when lineage decisions occur informs not only our understanding of zebrafish lateral line hair cell regenerate but also efforts to induce hair cell regeneration in mammals. Therefore, the high spatio-temporal resolution data presented here can serve to determine whether different injury paradigms lead to different transcriptional responses and provide a foundation for comparative regeneration studies across species and different regenerating organs.

## Results

To determine the stages for transcriptomic analyses, we analyzed time lapse recordings of regenerating hair cells (Figure 1F). The vast majority of hair cells are dead within 30min of addition of neomycin, and dead hair cell debris is cleared between 3-5h hours. Hair cells regenerate rapidly, with the first new pair detected within 3- 5 hours after neomycin exposure (Figure 1F, arrow; (Romero-Carvajal et al., 2015)). To characterize the transcriptional changes that underly these fast behavioral responses, we performed single cell RNA-Seq (scRNA-Seq) using the 10X Chromium platform during homeostasis (homeo) and six time points after 30 min neomycin exposure to kill hair cells (Figure 1F, homeo, 0 min, 30 min, 1h, 3h, 5h and 10h). We dissociated 5dpf *Tg(she:H2A-mCherry)* fish, in which all neuromast cells are labeled (Peloggia et al., 2021). After sequencing, we integrated cells from all time points using Seurat (v3.2.0) and plotted them in two-dimensional UMAP space (Figure 1G). Cluster analyses identified the same populations we previously described in homeostatic neuromasts (Figure 1H, Figures 1B-E; (Lush and Piotrowski, 2014)). Figure 1I and Figure S1A show the expression of selected cluster marker genes in UMAP space (Table S1). We did not observe any obvious loss or gain of a particular population during any of the regeneration time points (Figures S1B-C).

We show the results of the transcriptome analyses of all 6 time points after hair cell death using all cells in Figure 1- Figure 5, and present the results of pseudotime analyses using only central support cells and cells belonging to the hair cell lineage in Figure 6 – Figure 8. The results of our regeneration analysis can be queried using publicly available databases: the shiny app entitled ‘Neuromast Regeneration scRNA-Seq’ shows feature plots, violin plots, dot plots and heat maps of the entire data set, whereas the link within this shiny app called ‘Pseudotime Analysis’ shows feature plots, violin plots, dot plots and line graphs of central cells and hair cell lineage (Figures S2A-B, https://piotrowskilab.shinyapps.io/neuromast_regeneration_scRNA-Seq_pub_2021/). The genes present in both databases are listed in table S2. In addition, the data has been deposited to gEAR, a database that allows comparison of our data with other published data from other species (Figure S2C, https://umgear.org/p?l=e0d00950)(Orvis et al., 2021).

The transcriptional responses to hair cell death are rapid and cohorts of genes are robustly down and up-regulated immediately after hair cell death (0min time point, Figure 1J). These fast transcriptional changes continue throughout regeneration. UMAPs of individual cell types, in which cells are grouped according to the degree of similarity in their transcriptomes, show that cells at the 0min time point have a drastically different transcriptome than homeostatic cells and the 30min time point (Figures 1K-N). Cells collected at the 30min and 1h time points are more similar to each other but distinct from the 3h-10h time points. 5 and 10h cells are transcriptionally most similar to homeostatic cells. These groupings are not caused by batch effects as cells collected at different time points possess similar number of UMI count distribution (Figure S1D). The largest number of transcriptional changes occur in central, A/P, D/V and hair cell progenitor cells, whereas amplifying cells, hair cell progenitors, young and mature hair cells show a much smaller response (Figures 1K-N, Figures S1E-G). Mantle cells also show robust transcriptional changes, even though they do not proliferate in response to neomycin-induced hair cell death (Figure S1H; (Romero-Carvajal et al., 2015)). We observed the largest number of upregulated transcripts at 3h after hair cell death, whereas between 0min- 1h more genes are down-than upregulated.

In summary, our transcriptome analysis of closely spaced time points revealed that transcriptome changes occur rapidly in support cell populations immediately after killing hair cells. The different regeneration time points can be roughly grouped into three modules: 0min, 30min/1h and 3h/5h/10h. 10h cells begin to resemble homeostatic cells. To determine what transcriptional changes occur between the three modules we performed differential gene expression between various combinations of time points and selected genes that changed in several cell types or specifically in some cell populations (Table S3 (differentially expressed genes) and Table S4 (plotted genes shown in Figures).

### Module 1 (0min): Modulation of progenitor/stem cell maintenance/placodal and injury/inflammatory response genes

Progenitor/stem cell maintenance/placodal genes are rapidly turned off after neomycin treatment but are re-expressed between 3- 5h (Figure 2, Figure S3, Table S4). In contrast, injury/inflammatory genes are rapidly activated after hair cell death but are also sharply turned off again after 30min- 1h (Figure 3, Figure S4, Table S4).

**Figure 2.**
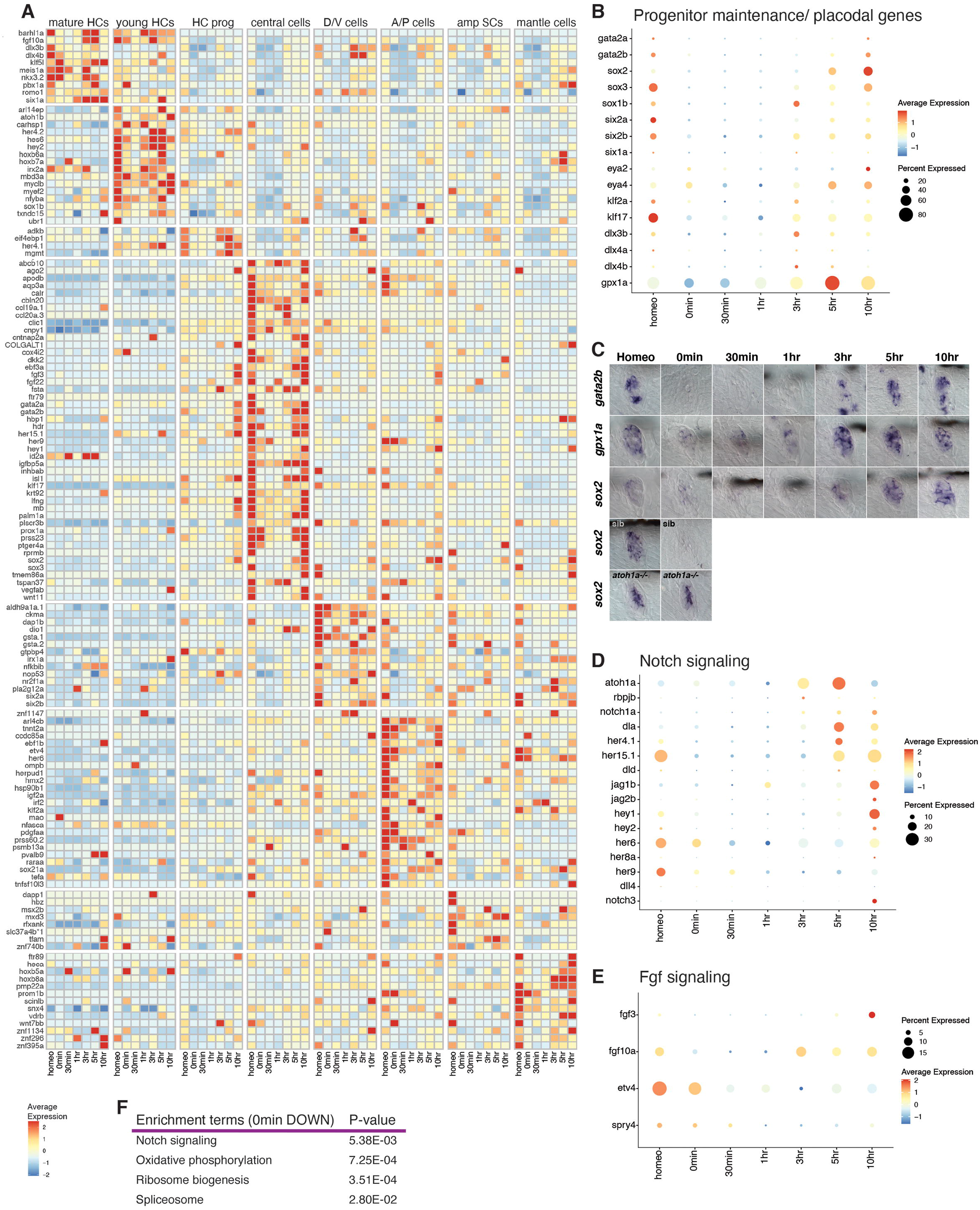
Downregulation of placodal genes, Notch and Fgf-signaling 0min after hair cell death. (A) Heatmap showing selected cell type-specific downregulated genes 0min after neomycin treatment. (B) Dot plot displaying the average expression of progenitor/stem cell maintenance/placodal genes during hair cell regeneration. The size of the dots represents the proportion of cells expressing the gene. (C) ISH images representing *gata2b*, *gpx1a*, and *sox2* expression during hair cell regeneration. Bottom two rows: ISH of *sox2* in sibling and *atoh1a* CRISPR mutant during homeostasis and 0min after neomycin treatment. (D and E) Dot plots showing the average expression of Notch (D) and Fgf (E) pathway members during hair cell regeneration. (F) Enrichment term analysis of downregulated genes 0min after hair cell death compared to all other time points.

**Figure 3.**
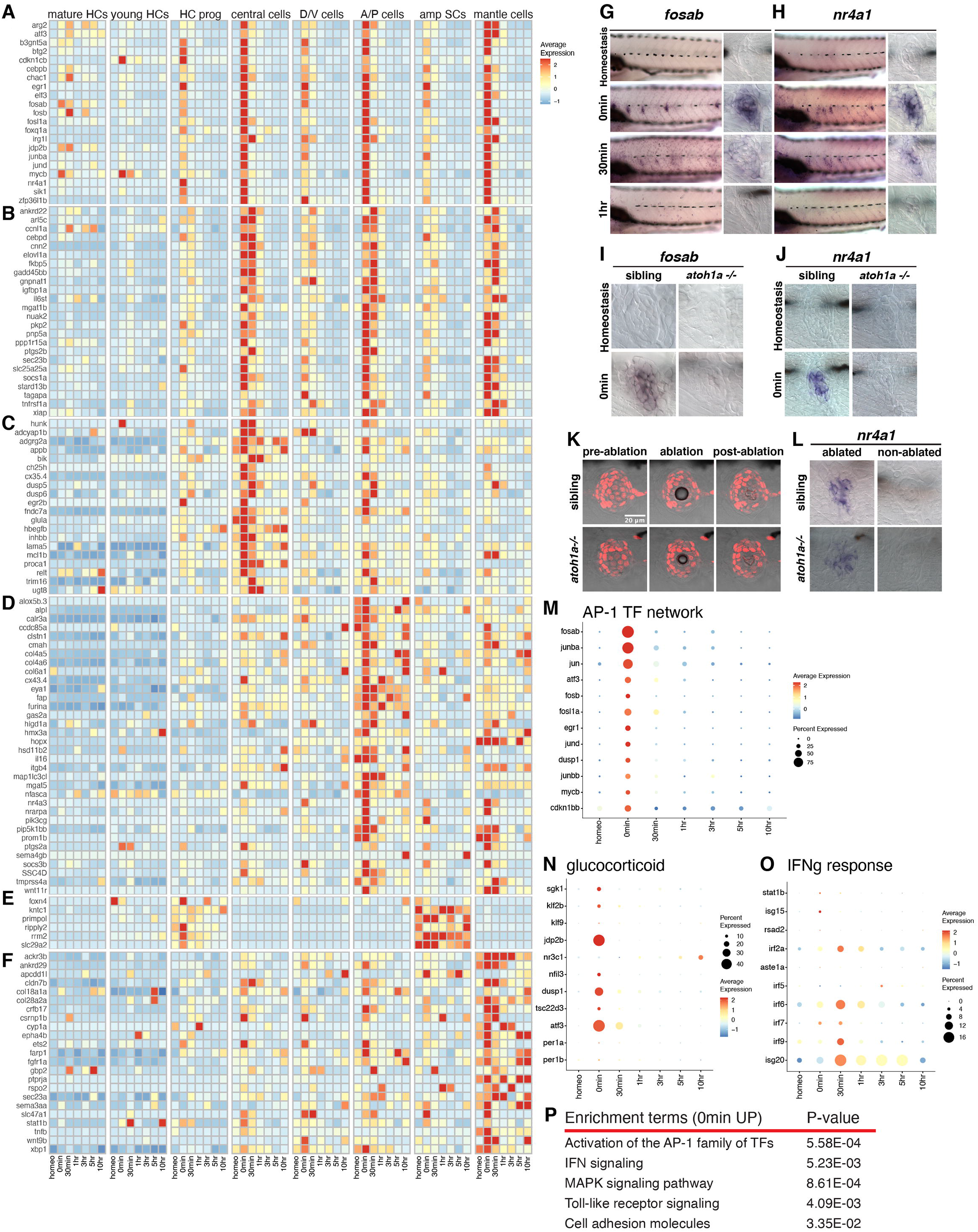
Injury/inflammatory response genes are rapidly upregulated 0min after hair cell death. (A - B) Heatmap of the expression of selected genes upregulated in all support cells at (A) 0min and (B) 0-30min. (C - F) Heatmaps of genes upregulated in different cell types at 0min and 30min after hair cell death. (G and H) ISH images of (G) *fosab* and (H) *nr4a1* in neuromasts at homeostasis, 0min, 30min, and 1h after hair cell death. (I and J) ISH images of (I) *fosab* and (J) *nr4a1* expression in neuromasts of sibling and *atoh1a* CRISPR mutant embryos at homeostasis and 0min after hair cell death. (K) Confocal images of *Tg(she:H2A-mCherry)* neuromast cells during laser ablation in 5dpf sibling and *atoh1a* mutant embryos. (L) ISH images of *nr4a1* expression in laser-ablated and non-ablated neuromasts of sibling and *atoh1a* CRISPR mutant embryos. (M - O) Dot plots displaying the relative expression of (M) AP-1 TF network members, (N) Glucocorticoid receptor signaling and (O) IFN signaling in neuromasts during hair cell regeneration. (P) Gene enrichment term analysis for genes upregulated 0min after hair cell death compared to all other time points.

#### Downregulation of progenitor/stem cell maintenance/placodal genes, Notch- and Fgf signaling 0min after hair cell death

0min after neomycin treatment, we observed that more genes are downregulated than upregulated (Figures 1K-N). A heatmap of the downregulated genes in different cell types and at different time points shows that the downregulation of genes at 0min is transient and that the majority of genes are re-expressed between 3-10h (Figure 2A, Table S4). It is also apparent that many downregulated genes are cell type specific. Among the downregulated genes are several involved in ear/lateral line placode development, including stem cell markers, such as *sox2/3*, as well as the Notch-, and Fgf signaling pathways (Figures 2B-E, Table S4; (Graham et al., 2003; Hernandez et al., 2007; Jiang et al., 2014; Sarkar and Hochedlinger, 2013)). We validated the downregulation of these genes by in situ analysis (Figure 2C, also (Jiang et al., 2014)). To test whether the observed downregulation is not caused by the neomycin treatment, we treated *atoh1a* mutant larvae, which lack hair cells with neomycin (Figure 2C). *sox2* is only downregulated after hair cell death in the siblings at 0min demonstrating that the downregulation of genes is not caused by the neomycin treatment but by the loss of hair cells. Figure S3A shows in which cell types different placodal genes are enriched. Prior studies showed that Notch and Fgf signaling are downregulated to allow for Wnt-dependent cell proliferation (Lush et al., 2019; Romero-Carvajal et al., 2015). Notch downregulation is also required to initiate hair cell differentiation (Ma et al., 2008; Romero-Carvajal et al., 2015; Wibowo et al., 2011). *vegfab* is also downregulated at 0 min but re-expressed by 10h (Figure 2A central cells). Interestingly, Vegf is required for hair cell regeneration in the avian ear (Wan et al., 2020), but a role for Vegf signaling in the lateral line has not been previously described.

Enrichment term analyses show that the most drastically downregulated processes are oxidative phosphorylation (OXPHOS), ribosomal proteins, genes involved in splicing and the Notch pathway (Figure 2F, Figure S3B, Table S3). These terms suggest that translation is inhibited and that the lateral line progenitor/stem cells undergo metabolic reprogramming from OXPHOS to glycolysis, a characteristic of other stem cells that become activated (Mathieu and Ruohola-Baker, 2017; Shyh-Chang and Ng, 2017).

#### Upregulation of the injury/inflammatory response at 0min: AP-1 complex, Tnf-, IFNg, Tgfb and glucocorticoid signaling

Injury either induces early response genes in all support cell types (Figures 3A-B) or only central-, A/P-, amp-, or mantle cells (Figures 3C-F, Table S4; (Bahrami and Drablos, 2016)). Some early response genes are known to be involved in an inflammatory response that, depending on the context and timing have either pro- or anti-regenerative effects (Eming et al., 2017; Kyritsis et al., 2012; Namdaran et al., 2012; White et al., 2017). We found that AP-1 complex members (eg., *fosab, fosb, fosl1a, jund, junba*), the nuclear hormone receptor *nr4a1* and other early response genes, such as *atf3*, *egr1*, and *btg2* are strikingly upregulated at 0min but are turned off again at 30min (Figures 3A, G, H, M, Figure S4A; (Cho et al., 2004)). Because AP-1 family members and *nr4a1* respond to injury/inflammation, we wondered whether the neomycin treatment, rather than hair cell death causes the upregulation of these genes. We treated sibling and *atoh1a* mutants that do not possess hair cells with neomycin and performed in situ hybridization with *fosab* and *nr4a1* (Figures 3I-J). *atoh1a* mutant neuromasts do not upregulate these genes in response to neomycin treatment demonstrating that they are specifically upregulated in response to hair cell death. We then tested whether *nr4a1* upregulation is induced by hair cell death specifically (Figures 3K-L). We laser-ablated support cells in siblings and *atoh1a* mutants and found *nr4a1* expression induced in both, demonstrating that death of any cell type leads to the upregulation of injury response genes.

Glucocorticoid and Jak/Stat signaling, as well as *immunoresponsive gene 1-like* (*irg1l)* are also activated in most cell types at 0min, but Jak/Stat and *irg1l* stay on until 30min/1h (Figure 3N, Figures S4A-B, D). *irg1l* is induced in keratinocytes by macrophages after infection and in turn leads to metabolic reprogramming in macrophages (Hall et al., 2014). Its role in lateral line regeneration needs to be determined. Interferon-, and Tgfβ signaling are activated slightly later than the early response genes and are expressed in distinct subset of cell types (Figure 3O, Figures S4C, D). Other genes are specifically enriched in particular cell types at 0min-1h, such as *egr2b*, *ch25h* and *hbegfb* in central cells and the autophagy gene *map1lc3cl* in AP cells (Figure 3A, Figures S4A, D). Why these genes are activated in some but not all cell types remains to be investigated. Other enrichment terms describing genes upregulated at 0min are MAPK signaling, Toll-like receptor signaling and cell adhesion molecules (Figure 3P, Figure S3B, Table S3).

### Module 2 (30min -1h): Regeneration-enriched genes are upregulated

30min- 1h genes are only lowly expressed in homeostatic sensory organs but are robustly induced for a short period during regeneration (Figure 4, Figure S5)

**Figure 4.**
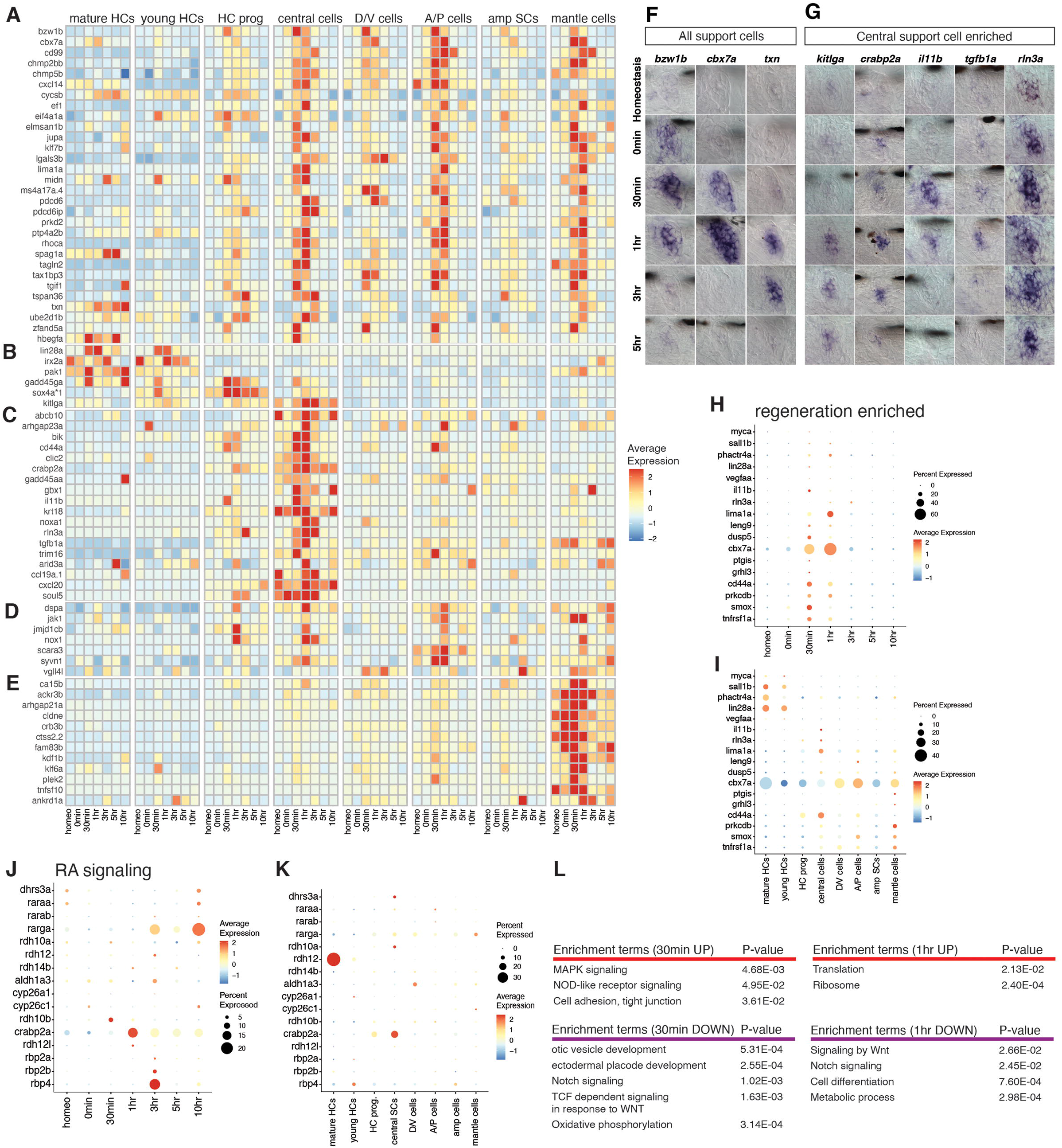
Upregulated genes 30min – 1h after hair cell death. (A) Heatmap of the average expression of selected upregulated genes in several support cell types and hair cell progenitors at 30min-1h. (B - E) Heatmaps showing the average expression of genes enriched in different cell types at 30min- 1h after hair cell death. (F and G) ISH images of genes upregulated at 30min-1hr in all neuromast cells (F) and central support cells (G). (H and I) Dot plots visualizing the average expression of (H) ‘regeneration enriched’ genes across seven different regeneration time points and (I) eight different neuromast cell types. (J and K) Dot plots of the average expression of RA pathway genes across (J) seven different regeneration time points and (K) eight different neuromast cell types. (L) Gene enrichment term analysis of up- and downregulated genes at 30min and 1h after hair cell death compared to all other time points.

#### Regeneration specific genes, translation, RA-, MAPK signaling and cell adhesion are induced between 30min- 1h

Module 2 genes are either upregulated in all support cells and hair cell progenitors at 30 min (Figures 4A, F, H, I, K, Table S4) or they are specifically upregulated in young and mature hair cells (*lin28a*, *irx2a* and *pak1*; Figures 4B, I); some central support cells and the hair cell lineage (*gadd45ga*, *sox4a1*, *kitlga*; Figure 4B); the central support cells (e.g. *cd44a*, *crabp2a*, *il11b*, *noxa1*, *rln3a*, *tgfb1a*, *arid3a*; Figures 4C, G); AP cells (Figure 4D) and mantle cells (Figure 4E). Retinoic acid (RA) signaling is also induced between 30min and 1h with pathway members being expressed in different cell types (Figures 4J-K, Figure S5). Enrichment terms that characterize upregulated genes at 30min are the MAPK cascade, NOD-like receptor signaling and Cell adhesion/tight junction, whereas ectodermal placode/otic vesicle development, Notch signaling, TCF-dependent Wnt signaling and oxidative phosphorylation are downregulated (Figure 4L, Figure S3B, Table S3). At 1h, ribosomal and other translation-associated genes are induced, whereas signaling by Wnt, Notch, cell differentiation and metabolism is still downregulated (Figure 4L, Figure S3B, Table S3).

### Module 3 (3h- 10h): Recapitulation of hair cell development

Genes expressed between 3h- 10h are expressed during development, as well as during hair cell turnover during homeostasis (Figure 5, Figure S6 and S7 (Romero-Carvajal et al., 2015)).

**Figure 5.**
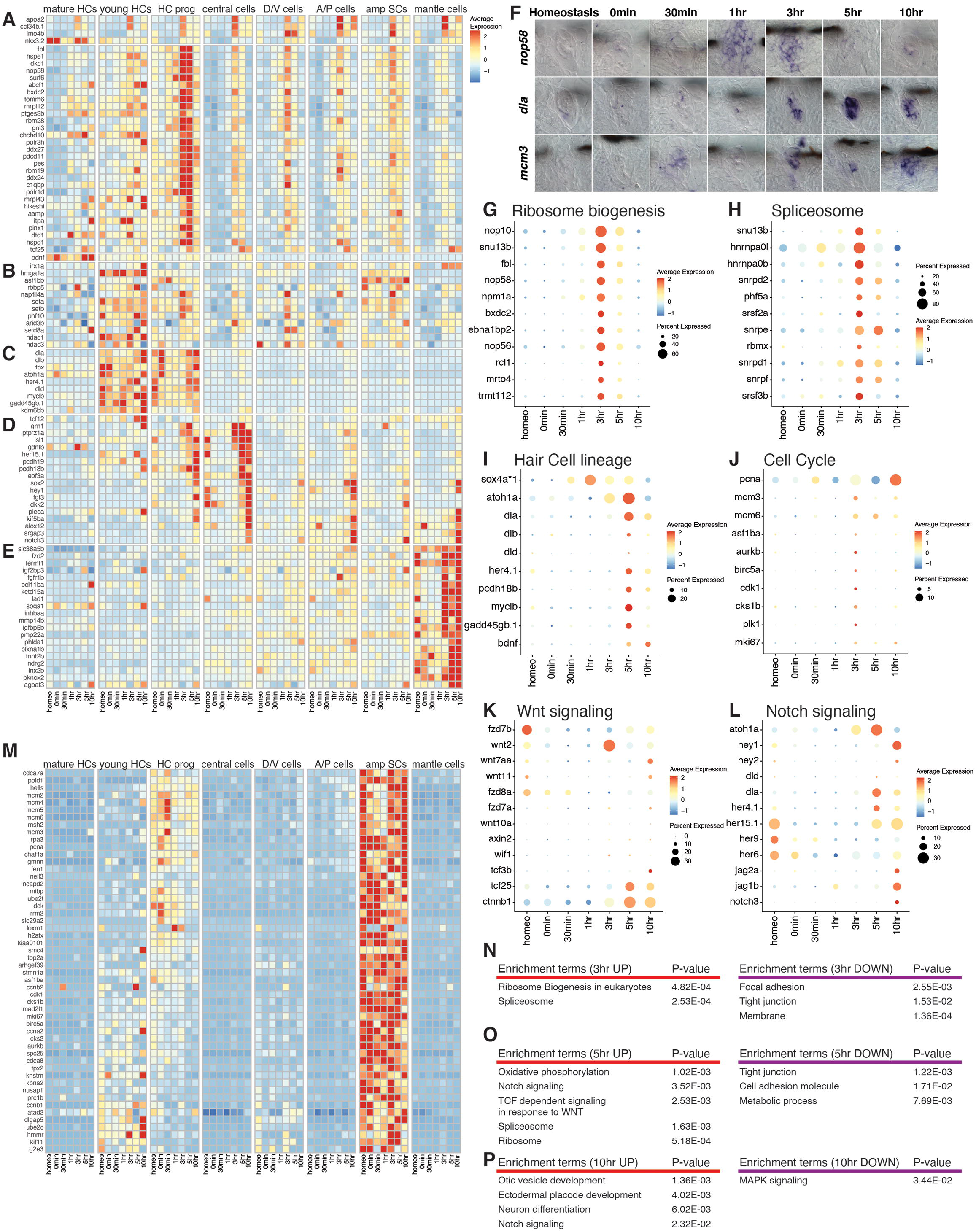
Metabolic reprogramming, ribosome biogenesis and hair cell lineage genes are enriched 3-10h after hair cell death. (A) Heatmap of the average expression of selected genes in several cell types at 3h- 10h after hair cell death. (B - E) Heatmaps of the average expression of cell type-specific genes upregulated 3h- 10h after hair cell death. (F) ISH images of *nop58*, *dla*, and *mcm3* in neuromasts during hair cell regeneration. (G - L) Dot plots displaying the average expression of (G) ribosome biogenesis, (H) spliceosome, (I) hair cell lineage, (J) cell cycle, (K) Wnt signaling, and (L) Notch signaling. The dot size shows the percentage of cells that express the gene. (M) Heatmap of the average expression of early and late cell cycle genes in eight different neuromast cell types during hair cell regeneration. (N - P) Gene enrichment term analysis up- and downregulated genes at (N) 3h, (O) 5h, and (P) 10h after hair cell death compared to all other time points.

#### At 3h the hair cell lineage becomes specified and ribosome biogenesis and splicing gene expression is activated

At 3h a few genes are upregulated in all cell types, such as *apoa2*, *ccl34b.1*, *lmo4b* and *nkx3.2* (Figure 5A, Table S4); however, the majority of 3h genes detected by 3h show cell-type specific expression (Figures 5B-E). Ribosome biogenesis genes and splicing factor expression particularly increase in HC progenitors, whereas epigenetic regulators are most highly expressed in young hair cells (Figures 5F, G, H, Figure S6A-B, Table S4). Figure S7 shows the heatmap of all genes shown in the dot plots, as not all are represented in Figures 5A-E. As a measure of the robustness of our scRNA-Seq, we generated dot plots and heat maps of ribosome biogenesis GO term genes and spliceosome pathway members listed in KEGG (Table S3). Indeed, these genes are highly enriched at 3-5h (Figures S6A-B). Thus, even genes that were not identified/filtered out in our differential gene expression analysis are expressed in our data set and can be queried using the shiny app or gEAR (Figure S2)(Orvis et al., 2021).

Also, the hair cell specifying gene *atoh1a* is upregulated in hair cell progenitors and young hair cells at 3h (Bermingham et al., 1999; Chen et al., 2002; Gubbels et al., 2008; Woods et al., 2004; Zheng and Gao, 2000), and downstream hair cell lineage targets, such as *delta* ligands, *myclb*, *gadd45gb*.1 and *kdm6bb* are upregulated by 5h (Figures 5C, I, Table S4). The percentage of hair cell progenitors and young hair cells that express these genes is shown in Figure S6C. In the heatmap Figures 5A-E, we omitted cell cycle genes. Figure 5J shows that cell cycle genes are also enriched at 3h and are most highly expressed in amplifying cells (Figure 5M, Table S4). Amplifying cells are located in the poles of the sensory organs and proliferate to give rise to more support cells (Figure 1D). They do not differentiate into hair cells unless they are displaced into the center of the sensory organ (Romero-Carvajal et al., 2015). Cell cycle genes are also activated in hair cell progenitors and young hair cells at 3h (Figures 5J, M). Early cell cycle genes involved in DNA replication, such as *mcm* genes, are enriched in hair cell progenitors and AP cells, whereas young hair cells and DV cells express genes characteristic of later stages of the cell cycle (Figure 5M). Thus, the hair cell lineage is tightly associated with proliferation in lateral line regeneration. Wnt signaling regulates proliferation in the lateral line (Jacques et al., 2013; Romero-Carvajal et al., 2015) and our data indicate that this pathway is reactivated at 3h (Figure 5K). We also find that Wnt pathway genes are enriched in the different support cell populations but are downregulated in hair cell progenitors and hair cells (Figure S6D).

Most notably, Notch pathway genes are re-expressed by 5h, as well as genes associated with oxidative phosphorylation (Figure 5L, Figures S6G-H) (Liberzon et al., 2015). Figures S6E-F show an expanded list of Notch pathway genes and the cell types in which they are enriched. We observe that ribosome biogenesis and splicing commence by 3h, whereas oxidative phosphorylation and the Notch/Wnt signaling pathways are reactivated by 5h (Figures 5N-P, Figure S3B, Table S3). Furthermore, focal adhesion, tight junction and cell adhesion genes are downregulated during these time points, a hallmark of proliferating epithelial cell. In fact, many of the genes associated with otic vesicle/ectodermal placode development such as *sox2* and *sox3* are re-expressed by 10h, while MAPK signaling and translation genes are markedly downregulated. Altogether, our data indicate that the discrete waves of gene up and downregulation observed during hair cell regeneration closely recapitulate key aspects of hair cell development.

### Pseudotime analysis indicates that central support cells either differentiate into hair cells or they re-express support cell genes

To gain a better understanding of the lineage trajectories that central support cells (central SCs) undergo during regeneration and to identify associated genes, we aligned central support-, progenitor- and hair cells along pseudotime using Monocle 3 (Cui et al., 2020; Qiu et al., 2017; Trapnell et al., 2014). We chose to limit the analysis to these cell types, as in vivo lineage tracing showed that central support cells are the direct precursors to hair cells (Lush et al., 2019). Although D/V and A/P cells can also give rise to hair cells (Romero-Carvajal et al., 2015; Thomas and Raible, 2019; Wibowo et al., 2011), it still needs to be determined whether A/P and D/V cells first acquire a central support cell fate. Therefore, we focused our analysis on central support cells to analyze hair cell and support cell lineage decisions.

To generate the UMAP in Figure 6A we specified 0min central cells (in purple) as the start of the trajectory. Cells with later pseudotime values are colored in yellow. The black line is the fitted trajectory along pseudotime and indicates lineage decisions that cells can take. In contrast to the differential gene expression performed on all cells from all time points (Figure 2- Figure 5), pseudotime analysis reveals a branch point at which cells either follow a path to hair cell differentiation (Figure 6B, lower branch) or they take a trajectory to revert back to a central support cell state (Figure 6B, upper branch). The pseudotime lineage trajectory model is temporally consistent with the regeneration time points that the cells were collected at, with 10h hair cells and 10h central support cells being at the end of each of the two lower and upper branch trajectories, respectively.

**Figure 6.**
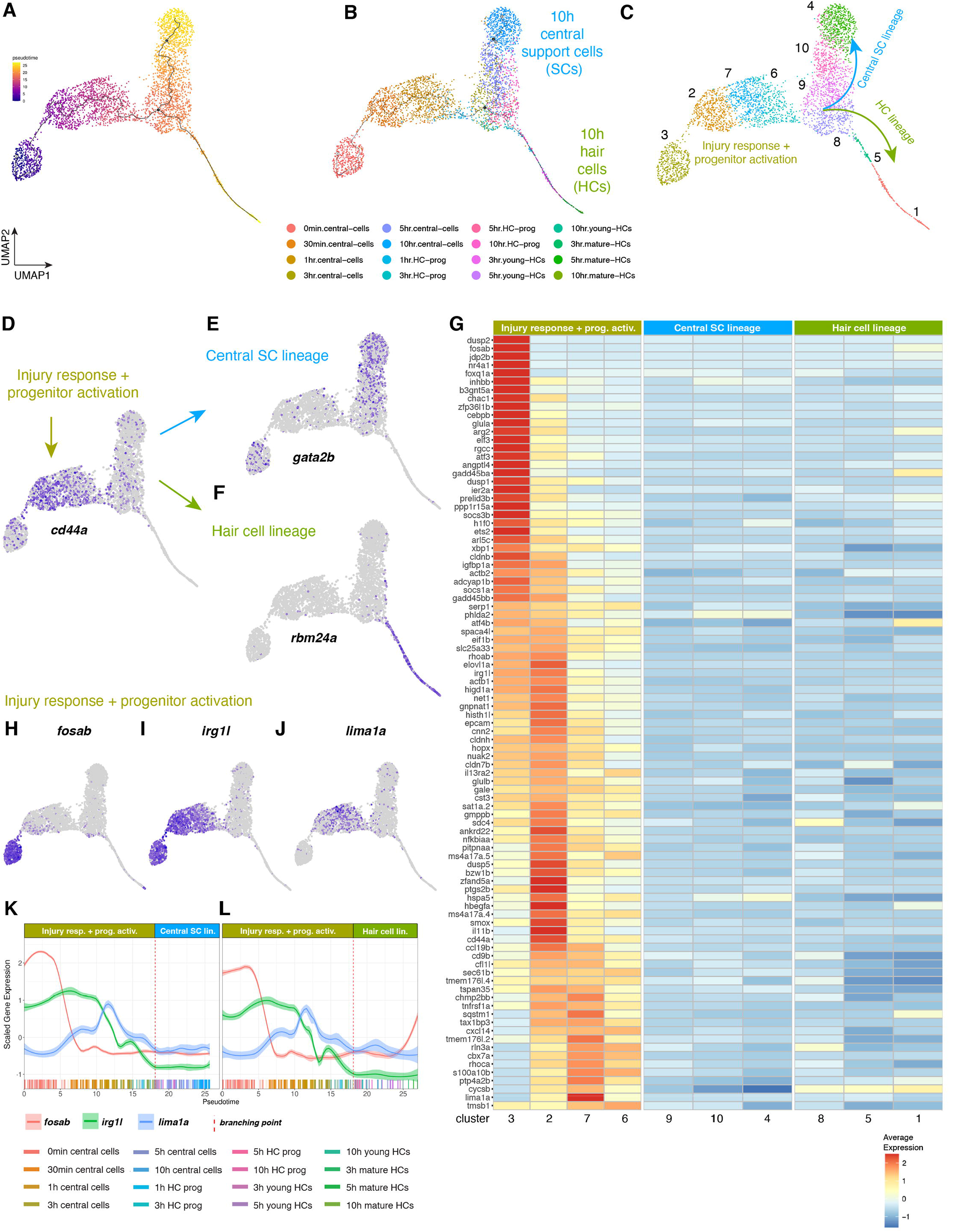
Pseudotime analysis (injury response + progenitor activation) (A - C) UMAPs of pseudotime trajectories of subsetted central support cells and hair cell lineage cells. (A) Dark purple shows cells at the beginning of pseudotime and yellow cells are at the end. (B) UMAP with the location of cells from different time points highlighted in different colors. (C) Cluster analysis groups cells with similar transcriptomes into ten different clusters and shows that, after the branch point cells either follow the central support cell (SC) lineage (blue arrow), or they follow the hair cell lineage (HC lineage, green arrow). (D) Feature plot of *cd44a* that is enriched in cells before the branch point. (E) Feature plot of *gata2b* that is enriched in the central support cell (SC) lineage. (F) Feature plot of *rbm24a* that labels the hair cell lineage. (G) Heatmap displaying markers selectively enriched in the injury response + progenitor activation population, partitioned into clusters. (H - J) Feature plots of genes expressed in the injury response + progenitor activation central support cell population. (K and L) Line plots showing the scaled expression of *fosab*, *irg1l*, and *lima1a* along the pseudotime trajectory in the (K) central support cell (SC) lineage and (L) hair cell lineage (HC). Individual cells are represented by color-coded tics on the x-axis.

The branch point at which cells are predicted to undergo a lineage decision occurs in cells at the 3- 5h time points (Figure 6B). To identify the genes that are expressed in cells before the branch point, we aggregated cells into Louvain clusters and performed a differential gene expression analysis between clusters of these different cell groupings/lineages (Figure 6C, Table S5). This analysis showed that sets of genes are progressively activated or de-activated in the injury response and progenitor activation cluster (Figure 6C clusters 3, 2, 7 and 6), the central support cell lineage (Figure 6C clusters 9, 10, and 4), and the hair cell lineage (Figure 6C clusters 8, 5, and 1). For instance, the injury response and progenitor activation clusters are characterized by the activation of *cd44a* (Figure 6D), while the central support cell lineage is marked by the upregulation of *gata2b* (Figure 6E). The hair cell lineage cluster is represented by the expression of *rbm24a*, an RNA-binding protein that regulates alternative splicing (Figure 6F).

#### Genes modulated in central support cells before the branch point

The heatmap (Figure 6G) and feature plots show genes that are immediately upregulated in cluster 3 (Figure 6C) central support cells in response to hair cell death but that are off in cluster 2, such as *fosab* (Figures 6 G, H, Figure S8A, Table S5); genes that are upregulated in clusters 3, 2, 7, 6, (Figure 6C) such as *irg1l* (consisting mostly of 30min and 1h central cells, Figures 6G, I, Figure S8A, Table S5); and genes that are most highly expressed in clusters 6 and 7 (Figure 6C) before the branch point, such as *lima1a* (Figures 6G, J, Figure S8A, Table S5). Line graphs of scaled gene expression of *fosab*, *irg1l* and *lima1a* in cells aligned along pseudotime clearly show these distinct gene expression dynamics (Figures 6K, L).

#### Genes modulated in the central support cell lineage after the branch point

To identify genes expressed in the central support cell lineage versus the hair cell lineage after the branch point, we specified the end and start points of each lineage and the algorithm assigned cells at and beyond the branch point to either lineage (Figure S8B). Cells before the branch point are shared between both lineages. The central support cell lineage chiefly consists of clusters 9, 10 and 4 (Figure 6C) and the expression dynamics of genes enriched in these clusters is shown in Figure 7A. Among the genes enriched in the central support cell lineage after the branch point are many of the placodal/stem cell-associated genes, such as *sox2*, *sox3*, *gata2a*, *gata2b*, *klf2a*, *klf17* and *prox1a* (Figure 7B; Figure S9, Table S5). *sox3* is already upregulated before the branch point but then rapidly turned off in the hair cell lineage (Figure 7B). Plotting the expression of these genes in a heatmap that contains all cell types and homeostatic cells shows that many of the central support cell lineage genes are also present during homeostasis. A number of these genes are also expressed in 10h A/P cells and mantle cells suggesting that these cell types also return to their homeostatic transcriptional state (Figure S9).

**Figure 7.**
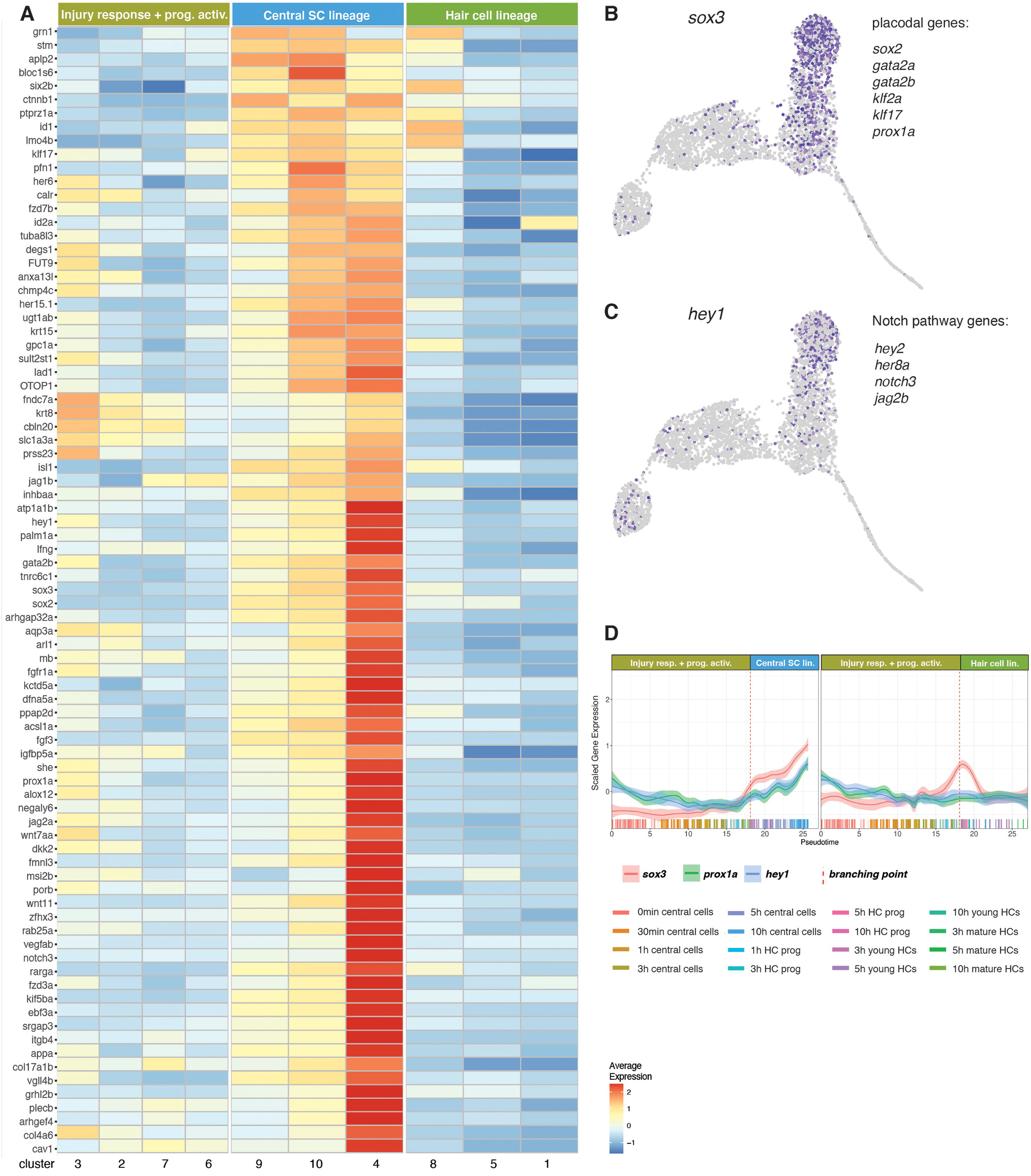
Pseudotime analysis (central support cell lineage after the branch point). (A) Heatmap of the average expression of genes that are selectively enriched in the central support cell lineage, partitioned by louvain clusters. (B) Feature plot of *sox3* and list of other genes expressed in these cells. (C) Feature plot of *hey1* and list of Notch pathway members that are also expressed in these cells. (D) Line plots of the scaled expression of *sox3*, *prox1a*, and *hey1* along the pseudotime trajectory in cells of the central support cell lineage and hair cell lineage cells (HC). Individual cells are represented by color-coded tics on the x-axis.

Notably, the Notch pathway is also upregulated (Figure 7C, Figure 5L) along pseudotime and highest in cluster 4 cells (Figures 6B-C, 10h central cells). Line plots of scaled gene expression along pseudotime show that *sox3*, *prox1a* and the Notch target *hey1* are upregulated in the central support cell lineage (Figure 7D).

We conclude from these analyses that the central support cells are on a trajectory back to becoming homeostatic central support cells. Indeed, the majority of the central support cell lineage genes are expressed at homeostasis but are subsequently downregulated at 0min, and are later re-expressed between 3-10h (Figure S9).

#### Genes modulated in the hair cell lineage

Atoh1 is required for hair cell specification but is not sufficient to regenerate a fully functional sensory epithelium when overexpressed in mammals of in culture (Burns and Stone, 2017; Jen et al., 2019). To identify additional genes that show similar expression dynamics as *atoh1a* or that could be acting upstream of *atoh1a*, we compared gene expression between central support cells and hair cell lineage clusters 8, 5 and 1 (Figure 6C, Table S5). Cluster 8 cells are hair cell progenitors and 5 and 1 consist of young and mature hair cells, respectively. Differential gene expression analysis between combinations of clusters shows the progression of gene expression along pseudotime (Figure 8A, cell cycle genes were excluded). We were particularly interested in identifying genes that might be involved in specifying hair cells and that act upstream of the hair cell specifier *atoh1a*. *sox4a* (called *sox4a*1* in our data sets) is an attractive candidate, as it is involved in hair cell development (Gnedeva and Hudspeth, 2015) and is expressed in a few central cells, as well as the hair cell lineage (Figure 8B; (Lush et al., 2019)). Indeed, our scRNA-Seq analysis shows that *sox4a* is upregulated before *atoh1a* at 30 min, whereas *atoh1a* is activated at 3h (Figure 5I, Figures 8B, C, Figures S10A-D). Both genes are expressed in central cells and the hair cell lineage (Figures S10C, D). We validated this difference in the onset of gene expression after hair cell death using HCR in situ hybridization (Figures 8E, F). Line plots of *sox4a*, *atoh1a* and *dld* show that *sox4a* is expressed before *atoh1a* and that *dld* is expressed after *atoh1a* is upregulated (Figures 8G, H).

**Figure 8.**
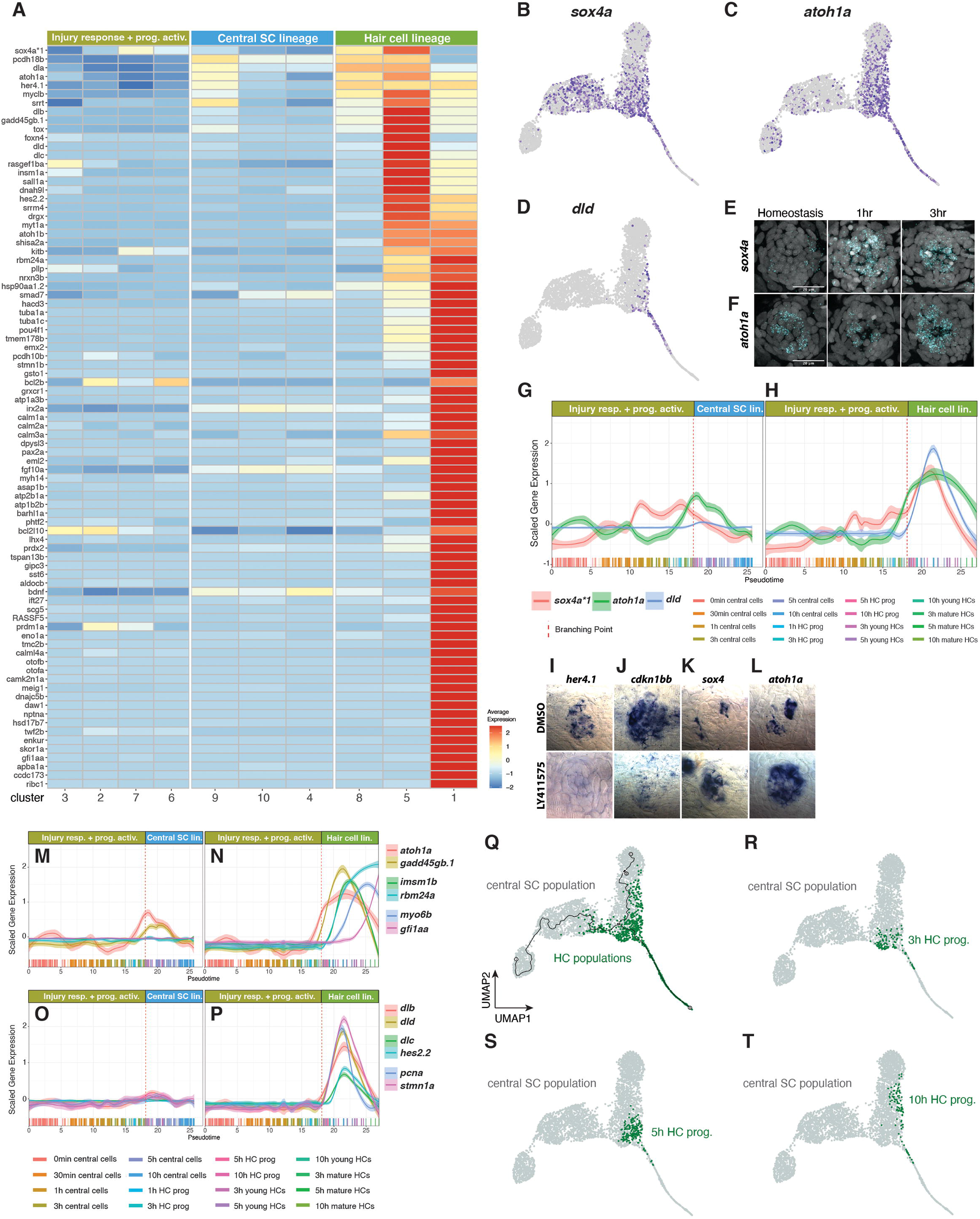
Pseudotime analysis (hair cell (HC) lineage after the branch point) (A) Heatmap of the average expression of genes that are selectively enriched in HC lineage cells, partitioned by louvain clusters. (B) Feature plot of *sox4a*. (C) Feature plot of *atoh1a*. (D) Feature plot of *dld*. (E) Fluorescent ISH (HCR) images of *sox4a* and *atoh1a* with DAPI in homeostasis, 1h and 3h after hair cell death. Scale bar: 20μm. (G and H) Line plots of *sox4a*, *atoh1a*, and *dld* scaled expression along the pseudotime trajectory in the (G) central support cell lineage and (H) HC lineage. Individual cells are represented by color-coded tics on the x-axis. (I - L) ISH images of *her4.1*, *cdkn1bb*, *sox4a*, and *atoh1a* in 5dpf DMSO- or LY411575-treated (to knock down Notch signaling) neuromasts. Larvae were treated for 6h before fixation. (M and N) Line plots of the scaled expression of *atoh1a*, *insm1b*, *myo6b*, *gadd45gb.*1, *rbm24a*, and *gfi1aa* along the pseudotime trajectory in the (M) the central support cell lineage and (N) HC lineage. Individual cells are represented by color-coded tics on the x-axis. (O and P) Line plots of *dlb*, *dld*, *dlc*, *hes2.2*, *pcna*, and *stmn1a* scaled expression along the pseudotime trajectory in the (O) central support cell lineage and (P) HC lineage. (Q) UMAP highlighting the distribution of HC populations comprising of HC prog, young HC and mature HCs. (R - T) UMAP showing the distribution of 3h, 5h, and 10h HC progenitor populations, respectively.

Notch signaling needs to be inhibited for hair cell specification genes to be expressed and we therefore wondered if *sox4a* is regulated by Notch signaling. Indeed, the downregulation of Notch signaling by the γ-secretase inhibitor LY411575, as evidenced by the loss of *her4.1* and *cdkn1bb* expression (Figures 8I, J) leads to the upregulation of *sox4a* and *atoh1a* (Figures 8K-L).

Line plots show temporal gene expression changes of well-known hair cell lineage genes with higher granularity than the heatmap in Figure 8A. Line plots reveal that the hair cell specifying genes *atoh1a* and the cell cycle regulator *gadd45gb.1* are upregulated in central support cells before the branch point (Figures 8M, N, stippled line), continue to increase in the hair cell lineage after the branch point (Figure 8N) but are downregulated in the central support cell lineage (Figure 8M). Within the hair cell lineage *atoh1a* and *gadd45gb.1* are followed by *insm1b*, *rbm24a*, *myo6b* and trailed by *gfi1aa* (Figure 8N). Line plots, the heatmap and feature plots also show that cell proliferation (*pcna*, *stmn1a*) occurs in the hair cell lineage (Figure 8P, *dlb*, *dld*, *dlc*, *hes2.2*) after the branch point and that cells in the central support cell lineage do not proliferate (Figure 8O, Figures S10E-F). The high-resolution characterization of gene expression dynamics in the hair cell lineage serves as a reference for developing protocols to induce hair cells in culture and to improve hair cells induction during mammalian regeneration.

### Hair cell progenitors likely revert back to a central cell fate

When we plotted the distribution of cells in the hair cell lineage along pseudotime, we observed hair cell progenitors clustering with the central support cell lineage (Figure 8Q, green cells). A more detailed analysis of hair cell progenitors from different time points showed that, whereas most of the 3 and 5h hair cell progenitors are located immediately after the branch point close to the hair cell lineage (Figures 8R, S), 10h hair cell progenitors group mostly with the central support cells (Figure 8T) suggesting that these cells had turned on hair cell genes but then failed to fully differentiate and instead turned on central cell genes. We hypothesize that initially more cells are competent to become hair cells but that the re-expression of Notch lateral inhibition signaling between 3-5h ensures that not all cells turn into hair cells but also into support cells (Figure S6E, Figure 7C; Romero-Carvajal., 2015). This interpretation is supported by the finding that only few of the *atoh1a* expressing central support cells later turn on *dld* (Figures 8C-D).

### Overview of cellular and molecular processes during the regeneration time course

To identify genes with similar trajectory-dependent expression dynamics and which may be co-regulated, we clustered these genes into units using Monocle 3 (Figure S11, Table S6). We identified eight gene units. To gain an understanding of the processes that characterize different stages of regeneration, we performed Enrichment term analysis on unit genes (Figure S11). Unit 3 is comprised of injury response genes. Units 8, 1, and 6 describe genes that are upregulated in cells before the lineage decision branch point and that are then enriched in the hair cell lineage, such as genes involved in ribosome biogenesis/translation, splicing and oxidative phosphorylation. Unit 2 contains genes of the central support cell lineage involved in sensory organ development and stem cell maintenance, and units 4, 7 and 5 contain hair cell lineage genes with the enrichment terms cell cycle, stereocilium organization, glycolysis and glutathione metabolism. Figure 9 synthesizes the dynamic changes of cellular and molecular processes that occur during the three modules of the hair cell regeneration process.

**Figure 9.**
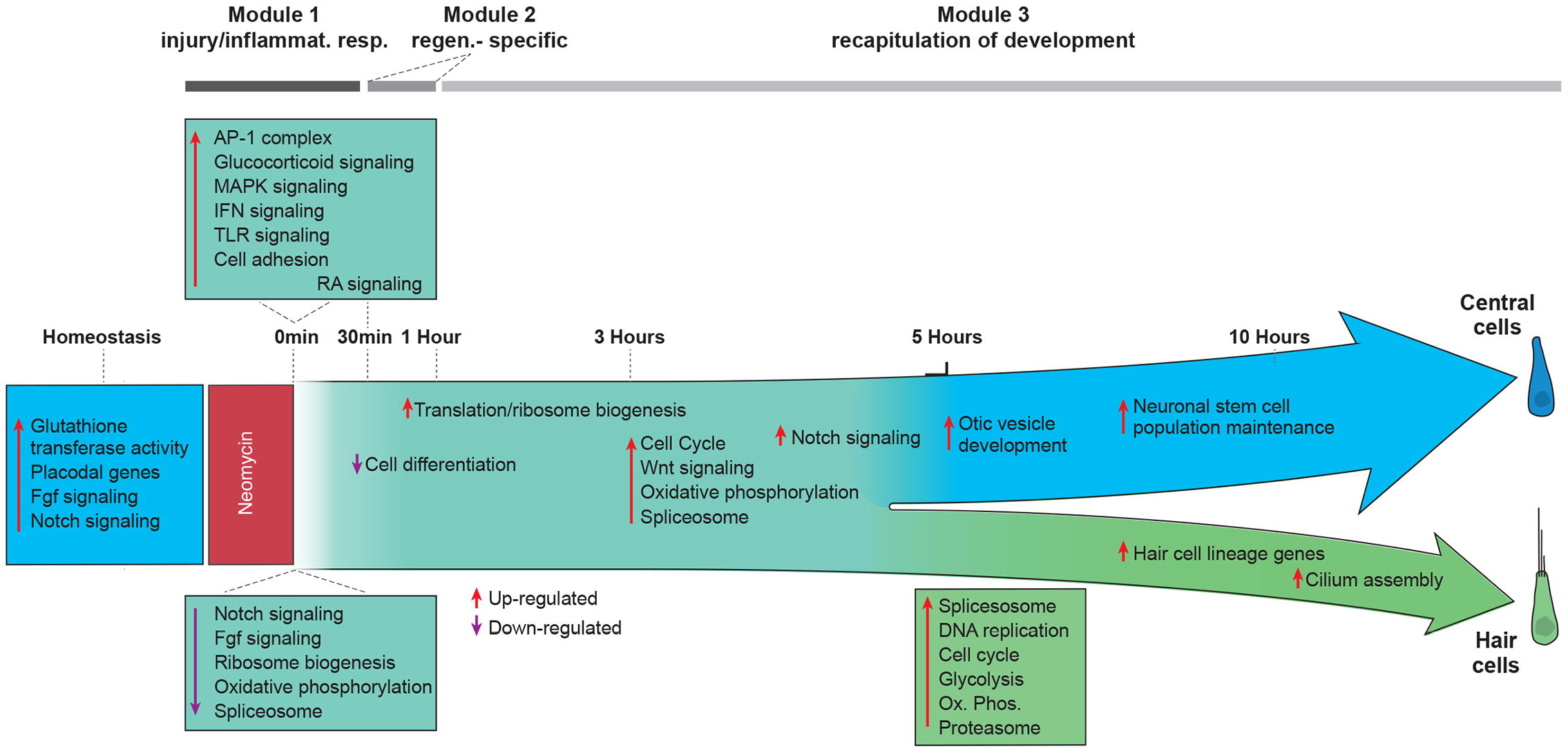
Overview of the molecular processes occurring during hair cell regeneration.

## Discussion

Inner ear hair cell and lateral line sensory organ development share key regulatory signaling pathways. Yet, unlike fish, mammalian hair cells do not regenerate once damaged. The remarkable speed (3- 5h) at which zebrafish regenerate lateral line hair cells after injury allowed us to characterize the gene expression dynamics of a regenerating vertebrate sensory organ at unprecedented resolution. Our results provide a blueprint of a gene regulatory network that enables support cells to re-enter the cell cycle and replace lost hair cells. Moreover, given that a number of genes/pathways involved in the regeneration of other sensory organs such as the retina in different species are also employed during lateral line regeneration, opens up the possibility that knowledge gained in a particular research organism or organ can inform studies in other species or organ systems (Denans et al., 2019; Hoang et al., 2020; Wan and Goldman, 2016).

### Gene expression changes occur within minutes after cell death

Our closely timed tissue collections demonstrated that drastic transcriptional changes occur within minutes after hair cell death. Recent studies showed that early response genes are required for triggering regeneration (discussed below). Yet, even though regeneration studies in different species increasingly include scRNA-Seq analysis, the majority of these studies collect only few time points and also collect the first sample many hours, or even days after injury. Our data indicate that in order to gain a complete understanding of the transcriptional dynamics underlying regenerative processes and to allow for comparative analyses between species, it is essential to characterize many closely spaced time points, starting within the first hour after injury.

### The injury/inflammatory response likely triggers regeneration

Genes expressed within minutes after hair cell death are injury or early response genes that are often involved in inflammation (Figure 3 and Figure 6G; (Bahrami and Drablos, 2016; Cho et al., 2004; Landen et al., 2016; Wang et al., 2020)). Inflammation has pro-regenerative effects in several contexts (Eming et al., 2017; Kyritsis et al., 2012; Namdaran et al., 2012; White et al., 2017). However, the timing of inflammation and its resolution by anti-inflammatory processes is crucial. For example, pharmacological suppression of the immune response before retina and CNS injury in zebrafish leads to a diminished regenerative response; however, blocking inflammation after injury speeds up the regeneration process (Kyritsis et al., 2012; White et al., 2017). These findings suggest that the inflammatory response facilitates the onset of the regeneration process but then needs to be shut off for regeneration to proceed.

We did not observe a strong pro-inflammatory response, such as robust *tnfa*, *il1b* or *il6* induction after hair cell death. However, *nr4a1*, a target of *il1b*, *tnfa* and *nfkb* is still strongly induced, as are AP-1 transcription factors (Figures 3A, M; (Pei et al., 2005)). Simultaneously, anti-inflammatory glucocorticoid signaling and *nfkb* inhibitors are rapidly activated at 0min (Figure 3). Interestingly, synthetic glucocorticoid agonists modestly enhance lateral line hair cell regeneration by increasing mitotic activity (Namdaran et al., 2012). Thus, it is possible that the short, pro-inflammatory signal observed in our study may be involved in facilitating hair cell regeneration by minimizing the detrimental effects of a long-term pro-inflammatory response (Denans et al., 2019; Landen et al., 2016; Monje et al., 2003). Jak/Stat3 is also induced at 0min. Loss of *stat3* function causes defects in hair cell regeneration/development in zebrafish and mice (Chen et al., 2017; Liang et al., 2012). The IFNG-pathway responds to hair cell loss at 30min (Figure 3O). It acts upstream of Jak/Stat1 and is involved in skeletal muscle regeneration. Yet, the role of the IFNG-pathway in hair cell regeneration remains to be investigated (Cheng et al., 2008; Zaidi, 2019). All of these early response pathways repress or activate each other in a context-dependent fashion that still needs to be characterized during hair cell regeneration (Cho et al., 2004; Xavier et al., 2016).

The AP-1 complex is of particular interest because it is not only required for wound healing but also for regeneration in several contexts (Beauchemin et al., 2015; Ishida et al., 2010; Raivich et al., 2004; Wenemoser et al., 2012). The AP-1 transcription factor complex consists of members of DNA binding proteins, such as Jun, Fos, ATF, JDP and Maf (Hess et al., 2004). Recent studies demonstrated that it is essential for triggering an injury and regeneration response by promoting chromatin accessibility (Beisaw et al., 2020). Additionally, AP-1-motif enriched regeneration responsive enhancers are likely a reason why non-mammalian vertebrates are better at regenerating (Wang et al., 2020). Even though mouse AP-1 enhancer elements drive gene expression in response to injury, they are unable to trigger a regenerative response, likely because of enhancer repurposing during evolution. It would therefore be important to test if AP-1 factors are required for hair cell regeneration in fish and if they are, compare the ability of the fish and mouse AP-1 motif containing enhancers to not only induce an injury- but also a regeneration response. Interestingly, noise induced hair cell damage in the rat cochlea leads to upregulation of the early response genes Egr1, Nr4a1(Nur77) and Btg2, but regeneration is not triggered supporting the notion of enhancer repurposing in mammals (Cho et al., 2004).

### Regeneration specific genes are active between 30min and 1h

Genes and pathways such as the retinoic acid pathway, which are enriched between 30min and 1h, are relatively regeneration specific, as they are not, or only lowly expressed during homeostasis. They show surprisingly short-lived activation and are again downregulated between 3- 5h. Many of these genes have been implicated in regeneration, such as *il11* that stimulates progenitor cells in axolotl tail regeneration, Jak/Stat3 and Nr4a1 target *relaxin3a (rln3a)* that is required for zebrafish heart regeneration (Figure 4G, (Fang et al., 2013; Tsujioka et al., 2017; You et al., 2018)). Likewise, the RNA binding protein Lin28 supports mouse axon regeneration, coordinates hair cell progenitor proliferation and differentiation and zebrafish hair cell regeneration (Doetzlhofer and Avraham, 2017; Wang et al., 2018; Ye et al., 2020). Lin28a also leads to enhanced tissue repair by metabolic reprogramming and increased glycolysis and oxidative phosphorylation in mice (Shyh-Chang et al., 2013). It is possible that these genes possess enhancers with AP-1 binding motifs and may represent the switch from a wound healing to a regenerative response, a hypothesis that can be readily tested in future work.

### Progenitor/stem cell maintenance/placodal genes are immediately downregulated

During the first 1h of hair cell regeneration more genes are downregulated than upregulated, which could represent a reprogramming step. A number of these genes, such as Notch/Fgf pathway members, *sox*, *six/eya, gata, dlx* and the *antioxidant enzyme glutathione peroxidase 1a* (*gpx1a)* are also expressed in the lateral line placode during development, and their homologs are involved in the development of hair cells in the mammalian inner ear (Figure 2, Figure S3A, (Basch et al., 2016; Kolla et al., 2020; Roccio et al., 2018)). Sox2/3, Six/Eya, Klf, Gata and Dlx transcription factors play multiple roles, such as stem cell maintenance, induction of pluripotency and lineage determination. *gpx1a* has been implicated in ES cell self-renewal (Wang et al., 2014). Likewise, we and others have previously shown that Notch and Fgf pathway downregulation is essential for support cell proliferation and that their re-expression at the end of the regeneration process is required for support cell maintenance (Lush et al., 2019; Ma et al., 2008; Romero-Carvajal et al., 2015). We propose that the downregulated genes are candidates that when temporarily inhibited might improve mammalian support cell reprogramming and hair cell regeneration (Iyer and Groves, 2021).

### Hair cell specification is triggered in the majority of 3- 5h central support cells but only a subset differentiates into hair cells

Between 3- 5h the majority of central support cells upregulate the hair cell specifier *atoh1a* (Figures 5I, 8C, 8F) but then only a subset also activate *dld* and enter the hair cell lineage (Figures 5I, 8D). The other cells maintain *atoh1a* expression for a few hours but simultaneously upregulate central support cell genes and enter the support cell lineage (Figures 8C, D). This finding is in line with what has been observed in Drosophila photoreceptor and zebrafish otic development where Atoh is first broadly expressed and acts as a proneural gene, but subsequently only a subset of the cells acquire a sensory fate (Cai and Groves, 2015). Unexpectedly, even a few *dld* expressing cells appear to revert back to the central support cell lineage (Figure 8D). Figure 8S-T show 5 and 10h hair cell lineage cells that align with central cells along pseudotime. These findings are consistent with recent reports that progenitors of different neural crest cell lineages initially express markers of both lineages and that lineage decisions arise over time (Soldatov et al., 2019). During hair cell development, the re-activation of Notch signaling at 3h correlates with the lineage bifurcation discovered in our pseudotime analysis (Figure 7C). We propose that the re-activation of Notch signaling in the central support cell lineage between 3- 5h may be responsible for ensuring that not all cells enter the hair cell lineage but that the majority of cells returns to a central support cell fate. This interpretation is supported by ours and our colleagues’ experimental data that prolonged Notch inhibition causes all cells to adopt a hair cell fate (Ma et al., 2008; Romero-Carvajal et al., 2015).

### Reactivation of Notch and Fgf signaling inhibits the cell cycle in the central support cell lineage

Interestingly, cell proliferation only occurs in the hair cell lineage and not in the central support cell lineage (Figures 8O, P, Figures S10E-F), which corelates with the re-activation of Notch and Fgf signaling in central support cells (Figure 7A). We previously showed that Notch signaling, and to a smaller degree Fgf signaling, inhibit proliferation via the inhibition of Wnt signaling during homeostasis (Lush et al., 2019; Romero-Carvajal et al., 2015). Thus, Notch does not only regulate lineage decisions but also inhibits proliferation and maintains cells in a progenitor state. The antiproliferative effect of Notch signaling is conserved in neural stem cells and the retina (Llorens-Bobadilla et al., 2015; Shin et al., 2015; Wan and Goldman, 2016). Likewise, Fgf signaling inhibits proliferation in the zebrafish retina (Wan and Goldman, 2017). Therefore, important unanswered questions are: what factors lead to the re-activation of Notch and Fgf signaling between 3-5h in the central support cell lineage? And what signals positively drive proliferation in hair cell progenitors upstream of Wnt signaling? (Jacques et al., 2013; Jacques et al., 2012; Romero-Carvajal et al., 2015; Shi et al., 2012; Wang et al., 2015).

Amplifying cells in the poles that do not directly give rise to hair cells also proliferate in a Wnt-dependent manner (Figures 1D, 5K heatmap; (Romero-Carvajal et al., 2015)) but it remains to be determined how their proliferation is inhibited during homeostasis, as Notch signaling is not active in the poles (Romero-Carvajal et al., 2015).

### Ribosome biogenesis and translation significantly increase between 3-5h

Concomitant with hair cell specification, the largest number of transcriptional changes occur at the 3h time point. Ribosome biogenesis and translation (e.g., *npm1a, nop56, nop58, RPL* and *RPS genes*) and splicing are activated particularly in the hair cell progenitors (Figures 5A, F-H, O, Figures S6A, B; (Lush et al., 2019)). An increase in ribosome biogenesis has also been observed in immature cardiomyocytes during heart regeneration in newborn mice (Cui et al., 2020). Similarly, ribosome heterogeneity is characteristic for stem cells and the role of regulated ribosome biogenesis during regeneration needs to be interrogated (Gabut et al., 2020; Li and Wang, 2020).

### Regeneration recapitulates later stages of hair cell development, but the trigger is regeneration specific

Many of the genes and pathways that we identified downstream of *atoh1a* in the hair cell lineage are involved in hair cell development in the mouse, chick and zebrafish implying that the hair cell specification and differentiation process is conserved between development and regeneration. The difference between regeneration and development is that regeneration is likely triggered by an injury/inflammatory response which fails to occur in mammals (Wang et al., 2020). Future studies need to focus on the injury/inflammatory response and downstream targets, to determine why these targets fail to be activated in mammals. Along this line, a gene expression atlas of mammalian inner ear sensory epithelia after damage is needed to identify the block in gene regulation that prevents mammals from regenerating.

### Similarities with zebrafish cardiac regeneration

Even though we have not yet performed a systematic comparison, we observed that many of the differentially expressed genes that we identified in the regenerating zebrafish lateral line have also been described in other regenerating organs, such as the retina, heart and fin (also (Denans et al., 2019)). For example, during zebrafish cardiac regeneration, the Jak1/Stat3 pathway members *jak1*, *jak3*, *Il6st* and *socs3b* are highly induced, as is *rln3a* (Fang et al., 2013). During zebrafish retina regeneration, several of the same growth factors/cytokines are induced, such as *hb-egf*, *insulin* and *il11* (Wan and Goldman, 2016). As in the lateral line, Notch downregulation leads to proliferation in the retina, which is enhanced by lin28a (Elsaeidi et al., 2018; Romero-Carvajal et al., 2015). Other genes involved in progenitor quiescence are *insm1a* and the Tgfβ inhibitor *tgif1a* (Chablais and Jazwinska, 2012; Lenkowski et al., 2013; Ramachandran et al., 2012). Likewise, some lateral line regeneration genes have also been described in fin and heart regeneration, such as *inhba/b* (Dogra et al., 2017; Sehring et al., 2016; Wang et al., 2020). This gene list is not comprehensive but stresses the value of comparing the regeneration programs between different organs and species to identify candidate regeneration genes. A recent comparison of the transcriptomes of fish, chick and mouse in Mueller glia after retina injury/regeneration revealed the strength of a comparative approach as they identified gene modules that are shared. However, it also uncovered difficulties. For example, TNF, MAPK, NFKB pathways are activated in mouse and chick Mueller glia cells after retinal injury (Hoang et al., 2020). However, these pathways were not identified in the zebrafish retina regeneration data set, as the earliest time point investigated was at 4h after retina injury, which based on our findings, is too late to observe the initial transcriptional responses. Thus, to gain a complete understanding of the transcriptional dynamics underlying regenerative processes and allow the comparison between species, it is essential to characterize many closely spaced time points, starting within the first hour after injury. High spatio-temporal resolution transcriptome studies from different regenerating species and organs are essential to test our hypothesis that the sequence of the three gene expression modules that we described is conserved and only the length of each module differs.

## Conclusion

We have generated a high-resolution regeneration scRNA-Seq data set that will serve as a foundation for devising strategies to help induce hair cell regeneration in mammals. The detailed transcriptome analyses reported here, combined with the characterization of three modules uncovered in our study will also serve as a reference to interrogate similarities in the regeneration response across different species/different organs and in response to different injury paradigms. Thus, this data is not only relevant for our understanding of hair cell regeneration but also for regenerative processes in general.

## Acknowledgments

We thank the Piotrowski lab members for insightful discussions and Drs. Alejandro Sánchez Alvarado, Nicolas Denans, Mark Lush, and Jeremy Sandler for critical reading of the manuscript. We thank Jillian Blanck, Jungeun Park, Allison Scott, Jeffrey Lange, Julia Peloggia for technical assistance, Drs. Joaquin Navajas Acedo and Mark Lush for in situ images, and Mark Miller for graphic design. We are also grateful to the Stowers Institute Core Facilities (Aquatics team, Cytometry Core, Imaging Core, Molecular Biology Core) for their technical expertise. We would also like to thank Dr. Ronna Hertzano, Joshua Orvis and Yang Song for developing and uploading our data to gEAR. This work was funded by an NIH (NIDCD) award 1R01DC015488-01A1 and by institutional support from the Stowers Institute for Medical Research to T.P.

## Author Contributions

S.B. designed and performed the experiments, analyzed and interpreted the data, and wrote the manuscript; N.T. analyzed and interpreted the data; D.D. analyzed and interpreted the data; Y-Y.T. performed experiments; J.N.A performed experiments; M.L performed experiments; T.P. designed the experiments, analyzed and interpreted the data, and wrote the manuscript.

## Declaration of interests

The authors declare no competing interests.

## STAR Methods

### RESOURCE AVAILABILITY

#### Lead contact

Further information and requests for resources and reagents should be directed to and will be fulfilled by the Lead Contact, Tatjana Piotrowski.

#### Data and code availability

Raw files such as BAM files and count matrices outputted by the CellRanger pipeline have been deposited in the Gene Expression Omnibus database, www.ncbi.nlm.nih.gov/geo (accession no. GEO will be submitted). Source code used to build our shiny applications are uploaded to the following repository, https://github.com/Piotrowski-Lab/shiny-apps. Supplementary scripts and wrapper functions used to process our seurat and pseudotime objects are available in the following github repositories: (https://github.com/Piotrowski-Lab/SeuratExtensions, https://github.com/Piotrowski-Lab/CellTrajectoryExtensions). All original source data are deposited in the Stowers Institute Original Data Repository and available online at https://www.stowers.org/research/publications/libpb-1636

### EXPERIMENTAL MODEL AND SUBJECT DETAILS

#### Zebrafish

Animal work followed the guidelines of the animal ethics committee (IACUC review board) at the Stowers Institute for Medical Research. The following zebrafish transgenic and mutant lines were used: *Tg(she:H2A-mCherry)^psi57Tg^*, *Tg(Myo6:H2B-mScarlet-I)^psi66Tg^* (Peloggia et al., 2021), *Et(Krt4:EGFP)^sqet4ET^*, *Et(Krt4:EGFP)^sqet20ET^* (Parinov et al., 2004), *atoh1a^psi69^* (Navajas Acedo et al., 2019).

### METHOD DETAILS

#### Sensory hair cell ablation

To ablate hair cells, 5dpf embryos were treated with 150∼300μM neomycin (Sigma-Aldrich, St Louis, MO, USA) for 30min at 28 °C. Following, embryos were washed with 0.5x E2 medium (7.5 mM NaCl, 0.25 mM KCl, 0.5 mM MgSO4, 75 mM KH2PO4, 25 mM Na2HPO4, 0.5 mM CaCl2, 0.5 mg/L NaHCO3, pH = 7.4) and incubated at 28 °C until further experimental needed.

#### scRNA-Seq

##### Embryo dissociation and FACS

5dpf larvae were anesthetized with tricaine for ∼1 min until they stopped moving (1:20 dilution of 4g/L tricaine in 0.5x E2 medium). We used ∼1000 mCherry-positive and 50 mCherry-negative (for gating control) larvae. To dissociate the larvae, we placed the larvae into four wells (250 larvae each) containing strainers (BD Falcon Cell Strainer (BD Biosciences, San Jose, USA), quickly rinsed the larvae in ice-cold DPBS and added 4.5 ml cold 0.25% trypsin-EDTA (Thermo Fisher Scientific, Waltham, USA). The larvae (250 each) were then transferred to one 15ml polypropylene conical tube (placed on ice) with a disposable transfer pipet. Then, the embryos were dissociated by trituration with a 1 ml pipette tip for 3min ∼ 3min 30 sec on ice. Since the neuromasts are superficial organs, we separated dissociated neuromast cells from the larval bodies by filtering the suspension through a Filcons 70 μm cell strainer (BD Biosciences, San Jose, CA, USA) into a 5ml polypropylene round-bottom tube. Subsequently, we centrifuged the cells at 2000 rpm (720 x g) for 7 min at 4°C. To wash off the trypsin-EDTA, we removed it, added ice-cold DPBS and centrifuged the cells at 2000 rpm (720 x g) for 7 min at 4°C. Resuspended cells in fresh ice-cold DPBS were filtered through a 35μm strainer (we only use strainer-cap part from 5ml blue cap strainer tubes (Falcon-Corning, Glendale, AZ, USA) into a falcon round bottom 5ml tube (snap cap tube, Falcon-Corning, Glendale, AZ, USA).

### FACS

To collect viable cells, we stained the resuspended cells with Draq5 and DAPI (Final concentration 25 μM DRAQ5 and 2 μg/ml DAPI). Following incubation for 5 min on ice, stained cells were sorted with BD influx cell sorter (BD Biosciences, San Jose, CA, USA) using a 100 μm nozzle at 20 psi and 1X DPBS (Sigma-Aldrich, St Louis, USA). Laser lines used are 355 nm, 488 nm, and 647 nm. DAPI, GFP, and DRAQ5 are collected using 460/50 nm, 528/28 nm, and 720/40 nm filters, respectively. FSC/SSC size selection gate was constructed by back-gating from high GFP expression. Targeted cells were collected into 90% MeOH (in DPBS) directly.

#### 10X Chromium scRNA-seq library construction

Methanol fixed cells were handled using the 10x Genomics suggested process and rehydrated with rehydration buffer (1% BSA and 0.5 U/μl RNase-inhibitor in ice-cold DPBS). The maximum recommended volume of single cell suspension (34 μl) was loaded on a Chromium Single Cell Controller (10x Genomics, Pleasanton, CA). Libraries were made using the Chromium Single Cell 3’ Library & Gel Bead Kit v2 (10x Genomics) according to manufacturer’s instructions. Following the library preparation, short fragment libraries were tested for quality and quantity using an Agilent 2100 Bioanalyzer and Invitrogen Qubit Fluorometer. The libraries were sequenced to produce a depth of ∼160-330M reads each on the Illumina HiSeq 2500 instrument (Illumina, San Diego, CA, USA) using Rapid SBS v2 chemistry with the following paired-end read lengths: 26 bp on the Read1, 8 bp on the I7 Index and 98 bp on the Read2.

#### scRNA-Seq read alignment and quantification

Raw reads were demultiplexed and aligned to version 10 of the zebrafish reference transcriptome (danRer10, Ensembl release 91) following 10X Genomics’s CellRanger (v2.1.1) pipeline. Prior to the downstream QC filtering we obtained the following cell numbers: 5,099 (homeostasis), 6,817 (0min), 6,727 (30min), 13,257 (1h), 3,314 (3h), 4,004 (5h) and 3,450 (10h) using CellRanger’s cell-association algorithm. Post-filtering, the number of cells per sample were 4335 (homeostasis), 2481 (0min), 2704 (30min), 2804 (1h), 2301 (3h), 1619 (5h), and 1829 (10h). The mean number of genes per sample were 726 (homeostasis), 653 (0min), 639 (30min), 650 (1h), 906 (3h) 826 (5h) and 797 (10h). All raw data for temporal samples including sorted BAM files and count matrices produced by CellRanger has been deposited in Gene Expression Omnibus (GEO) database, www.ncbi.nlm.nih.gov/geo (accession no. GEOx).

#### Pre-processing, quality filtering and batch integration

To distinguish zebrafish repeated gene symbols with unique Ensembl IDs, we modified the count matrices outputted from the CellRanger pipeline using a custom R script (https://github.com/Piotrowski-Lab/SeuratExtensions). Each repeated gene symbol is annotated with an asterisk followed by an incrementing number. A comprehensive gene list with unique repeated symbol annotations can be downloaded from our interactive web application (https://piotrowskilab.shinyapps.io/neuromast_regeneration_scRNAseq_pub_2021/). Low quality cells or cells containing doublets with reads greater than 6000 and less than 300 genes per cell after combining all samples were filtered from the subsequent analysis. Genes present in less than 10 cells were also removed from the dataset.

All seven temporal samples were integrated together following the standard integration pipeline outlined by the R package Seurat (v3.2.0, (Butler et al., 2018)). Here, individual temporal samples were normalized independently using default parameters via Seurat::NormalizeData. The log-normalized expression values are then z-scored on the integrated object via Seurat::ScaleData using default parameters after finding anchoring cells between samples.

#### Dimensional reduction, and cell classification

Choosing an optimal number of principal components (PCs) for dimensional reduction was determined by scree plot using Seurat::ElbowPlot. We selected PCs showing the greatest variance explained until each subsequent PC showed little to no change. We specified 50 total number of PCs to compute and selected the first 10 PCs based on the scree plot to build and shared nearest neighbor (SNN) graph using Seurat::RunPCA and Seurat::FindNeighbors, respectively. Seurat::FindClusters was used with a resolution of 0.4, resulting in 13 clusters. To visualize cells in two-dimensional latent space, we used UMAP dimensional reduction technique via Seurat::RunUMAP using the first 10 PCs aforementioned. Classification of cell clusters of the integrated dataset were annotated by calculating differential marker expression via Seurat::FindAllMarkers using default parameters. Annotating universal neuromast cell types preserved throughout regeneration was achieved via a comprehensive marker list previously generated by (Lush et al., 2019). Skin contaminant populations were removed from the subsequent analysis.

#### DE analysis or primary analysis

To distinguish differentially expressed marker genes between time points, we used Seurat::FindMarkers to compare cells in one query time point against all other cells (Figure 2-5, Table S3). The time point specific differential gene list was subsequently used in our gene set enrichment analysis. For all differentially expressed gene tables, we defined our statistical test using Wilcoxon Rank-test with default filtering parameters. Only genes with a p-value less than 0.05 were retained. The top 25 differential genes in each timepoint illustrate the expression dynamics between time points (Figure 1J).

#### Gene set enrichment analysis

Enriched terms at each time point were identified via the R package ClusterProfiler (v3.14.3, (Yu et al., 2012)) and ReactomePA (v1.30.0, (Yu and He, 2015)). Visualizing enriched terms was achieved via R package enrichplot (v1.6.1, https://github.com/YuLab-SMU/enrichplot). To explore terms from multiple pathway databases, we ran ClusterProfiler::gseGO, ClusterProfiler::gseKEGG, ReactomePA::gsePathway to access terms in the GO, KEGG, and Reactome databases respectively. To accommodate pathway databases that only read in Entrez IDs such as KEGG and Reactome, we converted Ensembl IDs from our differential gene tables to its complementary Entrez IDs using ClusterProfiler::bitr. For each time point analysis, we passed in a sorted named vector of mapping IDs and corresponding average log-fold change values to the enrichment functions aforementioned. We filtered and integrated meaningful terms, shown in Figure S3B, from the three pathway databases into a custom terms database via clusterProfiler::GSEA and returned a gseResult object. Here, we specified the minimal size of each gene set as 2, the maximum size of genes annotated for testing as 800 and 10,000 permutations. From the terms summary of each time point analysis, we labelled activated or suppressed terms by filtering positive and negative normalized enrichment scores (NES) respectively. All terms summaries from each time point analysis were then concatenated into a single dataframe to build the dotplot in Figure S3B (Table S3).

#### Pseudotime Analysis

The pseudotime analysis was processed using R package monocle3 v0.2.1(Cui et al., 2020; Qiu et al., 2017; Trapnell et al., 2014). From our integrated Seurat object, we subsetted the following hair cell populations: central cells at 0min through 10hr, hair cell progenitors at 1hr through 10hr, young hair cells at 3hr through 10hr and mature hair cells at 3hr through 10hr, totaling 3828 cells. Dimensional reduction was performed in UMAP latent space using Seurat’s standard preprocessing pipeline. We extracted the subsetted counts matrix and corresponding metadata and converted the processed Seurat object into a cds object. We coerced the UMAP embeddings previously generated in Seurat into its corresponding slot in the CDS object. A hallmark feature in the development of monocle3 is its scalability to partition cells to learn multiple, disjoint trajectories via partitioned approximate graph abstraction (PAGA). Since our subsetted cell populations have a common transcriptional ancestor, we set all possible partitions to a single partition to allow the trajectory path to travel through the entire UMAP embeddings. Thereafter, we fit a principal graph through the entire UMAP space via monocle3::learn_graph. Here, we controlled for the amount of branching paths by pruning trajectories that have a path diameter length less than 15. Next, cells were ordered along pseudotime via monocle3::order_cells, in which we specified the root of origin at the 0min central cell population. We relabelled cells belonging to the two major branching events as the Central Cell and Hair Cell Lineage via monocle3::choose_graph_segments, where we selected the start and end node pertaining to each lineage path. Cells that are shared between the two lineages prior to the branching point were relabelled as the Injury Response + Progenitor Activation Lineage. To retrieve the exact pseudotime value and UMAP embeddings of the branching point, we built a custom wrapper function, CellTrajectoryExtension::get_branching_point, found in our github repository (https://github.com/Piotrowski-Lab/CellTrajectoryExtensions). In brief, we built a dataframe that includes every node label with its corresponding UMAP embeddings and subsetted for branching nodes using monocle3::branch_nodes. We then identify the approximate cell barcode and corresponding pseudotime value pertaining to the branching node’s embeddings.

To identify correlated gene expression along the trajectory, we utilized a spatial autocorrelation test statistic implemented in monocle3::graph_test, which returns a Moran’s I value for each gene. To retain all positive, statistically significant autocorrelation values from monocle3::graph_test, we subsetted gene entries with Moran’s I greater than or equal to 0.10 and q-value less than 0.05. We then passed in our filtered genes from the graph_test output into monocle3::find_gene_modules to cluster genes into meaningful modules (called units in the text) using Louvain clustering. Here, we parameterized our resolution at 0.0005 and used the first 3 PCs to cluster our genes, resulting in 8 units as shown in Figure S11 To illustrate dynamic expression of genes in each unit, we calculated the z-scored expression of each gene in its corresponding module and averaged the expression of the cells along pseudotime. We then fit a smoothing line connecting the relationship between average expression of genes in each unit and pseudotime using a locally weighted smoothing method (LOESS). We set our degree of smoothing to 20% using the span argument.

#### Interactive web application and data repository

To encourage fast and dynamic visualization of our scRNA-seq data, we built an interactive web-based graphical user interface (GUI) via R base Shiny (https://piotrowskilab.shinyapps.io/neuromast_regeneration_scRNAseq_pub_2021/). The datasets imported in the shiny app includes the integrated neuromast regeneration and the pseudotime analysis. Features include visualizing user inputted gene expression via featureplots, dotplots, heatmaps, pseudotime line plots and comparative differential analysis. Instructions are provided on the homepage of the web app. In addition, our integrated regeneration data is uploaded to the Gene Expression Analysis Resource (gEAR) website (https://umgear.org/p?l=e0d00950)(Orvis et al., 2021).

#### Confocal Imaging and laser ablation

For confocal imaging, zebrafish embryos were embedded in 1% low-melt agarose (Promega, Madison, USA), which contains 1x tricaine in 0.5x E2 medium in glass bottom 35 mm dishes (MatTek, Ashland, USA). Imaging was performed using a Zeiss LSM 780 confocal microscope with 10x or 40x objectives. All images were processed using Fiji (Schindelin et al., 2012) software. To ablate specific neuromast cells, 5dpf of *atoh1a* sibling or mutant embryos *Tg(she:H2A-mCherry)^psi57Tg^* were mounted in 1% low-melt agarose with 1x tricaine in 0.5x E2 medium in glass bottom 35 mm dishes. Neuromasts were identified and ablated using an LSM-780 (Zeiss) confocal microscope. Tissue fluorescence was excited at 561nm through a 40x LD C-Apochromat objective (Zeiss, NA = 1.1) and collected back through the same objective, then through a 562-633nm bandpass filter. Photons were detected with a GaAsP detector running in photon counting mode. Once the tissue of interest was identified, ablation occurred with a Ti Sapphire 2 photon laser (Coherent) running at 800nm in mode-locked mode. The laser power at the back aperture of the objective was 6mW. Ablation occurred over a user selected ROI by point scanning the 2 photon beam through the ROI a user specified number of times. These settings were adjusted as needed to achieve ablation. Successful ablation was verified by imaging the tissue in transmitted light.

#### Notch inhibition

5dpf wild type larvae were treated with 50μM LY411575 (Selleckchem, Houston, TX, USA) or 1% DMSO (Sigma-Aldrich, St Louis, MO, USA) as a negative control for 6 hours, then fixed overnight in 4% paraformaldehyde at 4°C.

#### In Situ Hybridization

For conventional ISH, anti-sense RNA probes were generated by PCR with cDNA from mixed stages of zebrafish embryos using T3 promoter-tagged reverse primer (Table S7), except *gata2b* (Butko et al., 2015), *sox2* (Kudoh et al., 2001), *txn, cdk1nn, her4.1* (Jiang et al., 2014), *dla* (Appel and Eisen, 1998), *atoh1a* (Itoh and Chitnis, 2001). Go-Taq (Promega, Madison, USA) was used for PCR reaction and PCR product was purified by NEB gel extraction kit (New England Biolabs, MA, USA). Following that, In vitro transcription was performed with T3 RNA polymerase (Promega, Madison, WI, USA) and DIG-labelled kit (Roche, USA) in a PCR machine at 37°C for 4 hrs. After DNase I treatment at 37°C for 30mins, In vitro transcribed RNA was purified using RNA clean and concentrator kit (Zymo Research, Irvine, CA, USA) in accordance with the manufacturer’s manual. DIG-labelled RNA probe was diluted as 10 ng/μl in hybridization buffer and stored at −20°C. Colorimetric in situ hybridization was performed as described previously (Navajas Acedo et al., 2019). Hybridization chain reaction (HCR) for *sox4a*-B2 and *atoh1a*-B2 (Choi et al., 2018) was carried out following the manufacturer’s manual, except that we incubated in the probe and amplifier for 48h each. (Molecular Instruments). The amplifier used was B2-546 (Molecular Instruments). Stained and fixed larvae were stained with DAPI (5μg/ml) for 30 min at room temperature and washed three times with 5x SSCT and imaged with a confocal microscope.

## Supplementary Table Legends

**Table S1. Markers for the different cell type clusters.**

This spreadsheet contains the differentially expressed gene list (cluster markers) of neuromast cell types from the integrated regeneration dataset via Seurat::FindAllMarkers with default parameters. Genes with p-values greater than 0.05 were omitted from the gene list.

**Table S2. All genes present in the scRNA-Seq data set.**

This spreadsheet contains a list of all genes within the integrated neuromast regeneration scRNA-Seq dataset. Genes present in the expression matrix were extracted and placed within the gene table column named “Gene.name.uniq” along with their corresponding Ensembl ID, ZFIN ID, description, etc.

**Table S3. Differential gene expression between time points.**

This spreadsheet corresponds to the differential pseudobulk time point expression analysis. For each differential expression analysis, we compared cells labelling a query timepoint against cells in all other time points using the R call function Seurat::FindMarkers followed by a gene set enrichment analysis using the R package ClusterProfiler.

**Table S4. Genes represented in figures.**

This spreadsheet contains the list of genes used to plot heatmaps and dotplots corresponding to Figures 1-8, S1-S10, and partitioned into individual tabs. Each tab’s gene table contains genes symbols, corresponding Ensembl IDs, descriptions, as well as columns of the average scaled (normalized and z-scored) expression value grouped by the cell identities displayed on the x-axis.

**Table S5. Markers for clusters in the pseudotime UMAP.**

This spreadsheet contains lists of differentially expressed genes partitioned by clusters ordered along the pseudotime trajectory. We grouped clusters based on where they fall before and after the major branching event. We labelled clusters 3,2,7,6, which aggregate before the branching point as “Injury reponse + progenitor activation”. Clusters 4,9,10 with the end terminus leading to 10hr central cells, we labelled as “Central support cell lineage”. Clusters 1,5,8 with the end terminus leading to 10hr central cells, we labelled as “HC lineage”.

**Table S6. Genes belonging to different modules (units).**

This spreadsheet contains outputs of genes clustered into modules (units) along the pseudotime trajectory. Gene tables were generated via monocle3::find_gene_modules after subsetting for genes from the spatial autocorrelation analysis (monocle3::graph_test) with a moran_I greater than or equal to 0.10 and q-value less than 0.05.

**Table S7. Primers for in situ hybridization probe generation.**

## Notes

https://piotrowskilab.shinyapps.io/neuromast_regeneration_scRNA-Seq_pub_2021/

https://umgear.org/p?l=e0d00950

### Competing Interest Statement

The authors have declared no competing interest.

### Summary of Updates

The author list is updated.

