## Supplementary Figures for "High-resolution single cell transcriptome analysis of zebrafish sensory hair cell regeneration"

#### Figure S1

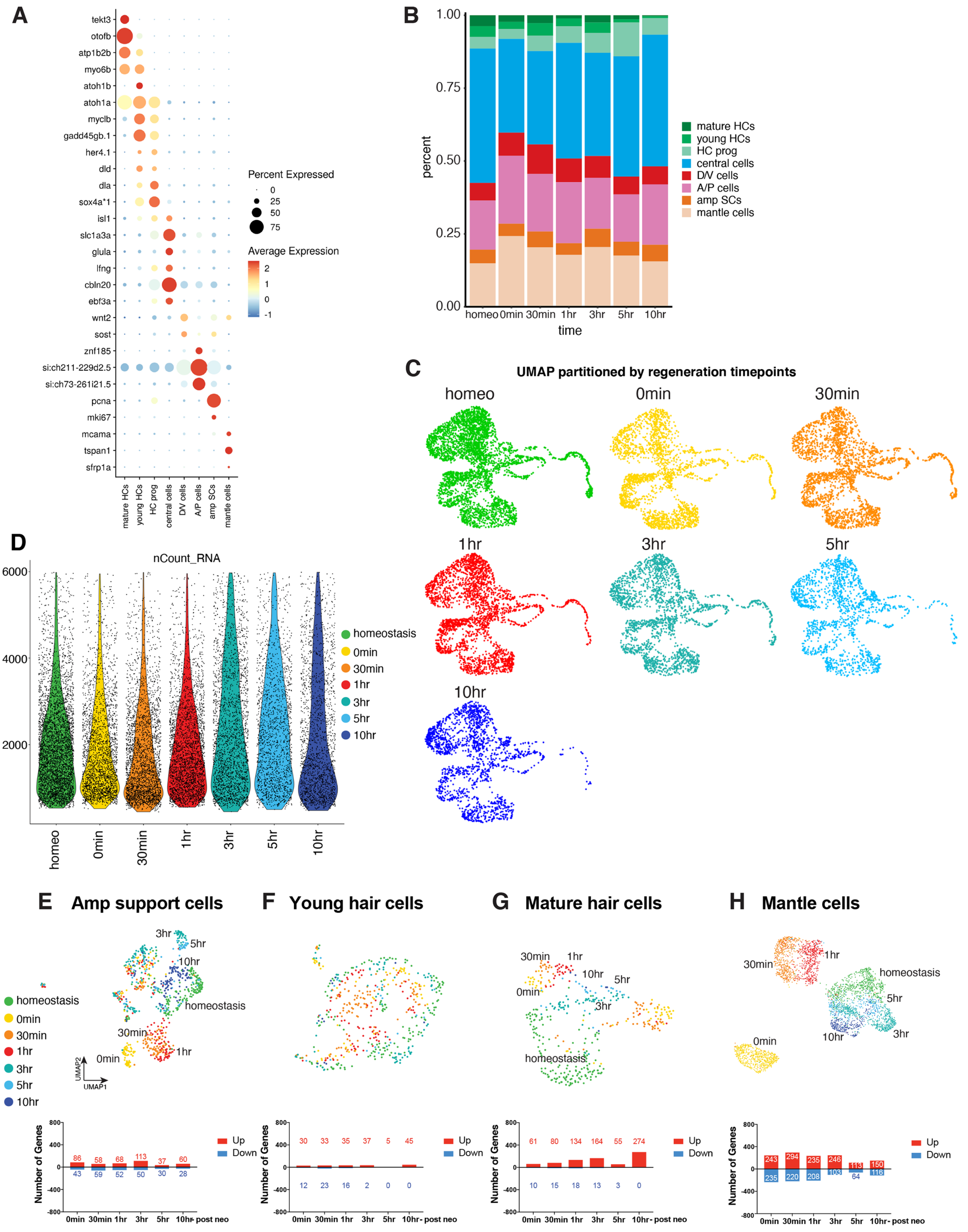
A
tekt3
otofb
atp1b2b
myo6b
atoh1b
atoh1a
myclb
gadd45gb.1
her4.1
dld
dla
sox4a*1
isl1
slc1a3a
glula
lfng
cbln20
ebf3a
wnt2
sost
znf185
si:ch211-229d2.5
si:ch73-261i21.5
pcna
mki67
mcama
tspan1
sfrp1a
mature HCs
young HCs
HC prog
central cells
D/V cells
A/P cells
amp SCs
mantle cells
Percent Expressed
0
25
50
75
Average Expression
2
1
0
-1
B
1.00
0.75
0.50
0.25
0.00
percent
homeo
0min
30min
1hr
3hr
5hr
10hr
time
mature HCs
young HCs
HC prog
central cells
D/V cells
A/P cells
amp SCs
mantle cells
C
UMAP partitioned by regeneration timepoints
homeo
0min
30min
1hr
3hr
5hr
10hr
D
nCount_RNA
6000
4000
2000
homeostasis
0min
30min
1hr
3hr
5hr
10hr
homeo
0min
30min
1hr
3hr
5hr
10hr
E
Amp support cells
3hr
5hr
10hr
homeostasis
30min
1hr
0min
UMAP2
UMAP1
homeostasis
0min
30min
1hr
3hr
5hr
10hr
F
Young hair cells
G
Mature hair cells
30min
1hr
10hr
5hr
0min
3hr
homeostasis
H
Mantle cells
30min
1hr
homeostasis
5hr
10hr
3hr
0min
Number of Genes
800
400
0
-400
-800
86 58 68 113 37 60
43 59 52 50 30 28
0min 30min 1hr 3hr 5hr 10hr- post neo
Up
Down
30 33 35 37 5 45
12 23 16 2 0 0
61 80 134 164 55 274
10 15 18 13 3 0
243 294 235 246 113 150
235 220 208 103 64 116

**Figure S1. Related to Figure 1.**

(A) Dot plot showing marker genes for eight different cell types.

(B) Proportion of cell types in samples of each time point.

(C) UMAPs showing the distribution of cells partitioned by individual time point post-integration.

(D) Violin plot visualizing the distribution of raw counts/cell by time point.

(E - H) UMAPs of (E) amplifying cells, (F) young hair cells, (G) mature hair cells, and (H) mantle cells with bar graphs showing the distribution of cells and the number of up and down-regulated genes per time point compared to homeostasis.

#### Figure S2

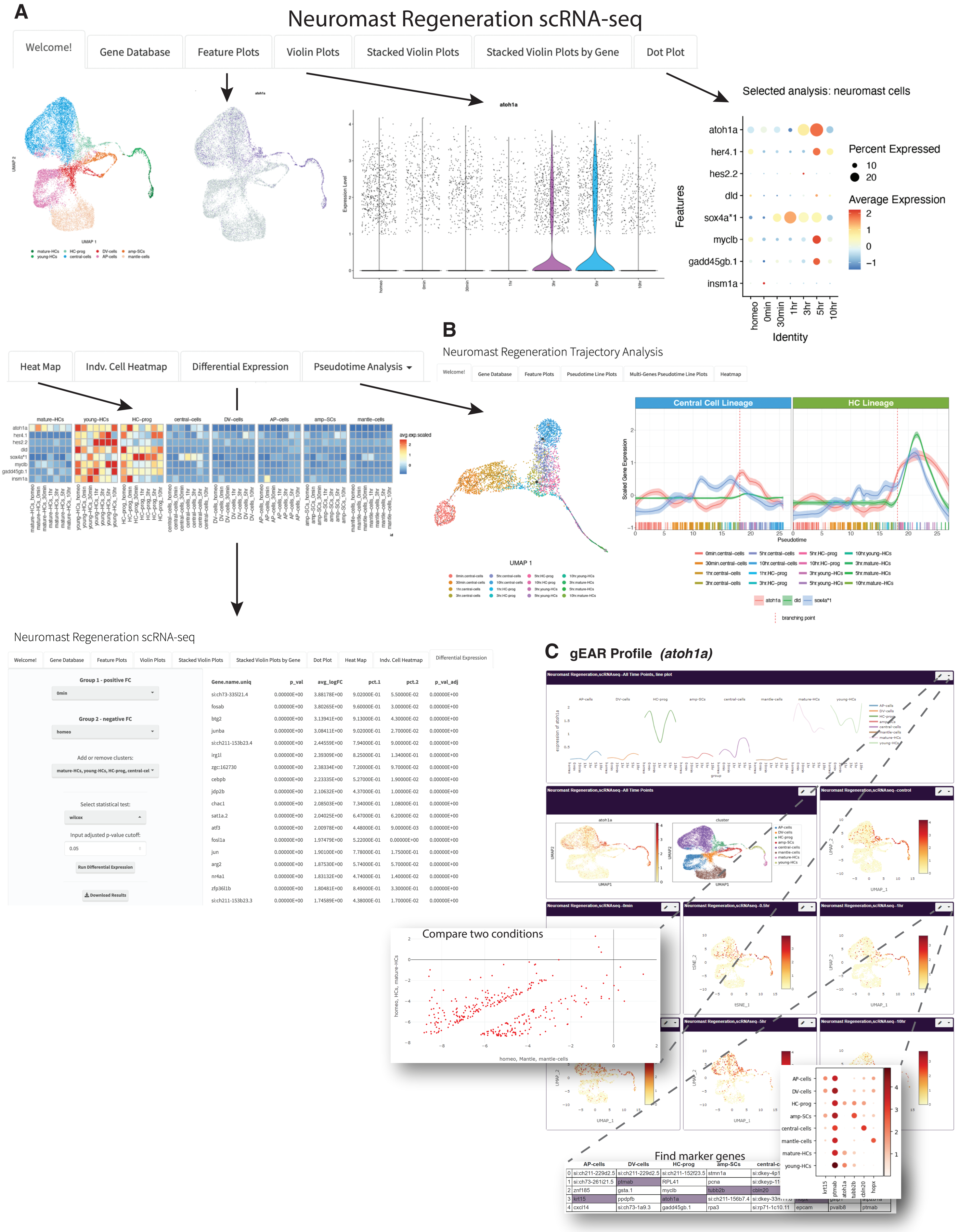
A
Neuromast Regeneration scRNA-seq
Welcome!
Gene Database
Feature Plots
Violin Plots
Stacked Violin Plots
Stacked Violin Plots by Gene
Dot Plot
atoh1a
Expression Level
homeo
0min
30min
1hr
3hr
5hr
10hr
Selected analysis: neuromast cells
atoh1a
her4.1
hes2.2
dld
sox4a*1
myclb
gadd45gb.1
insm1a
Features
Identity
Percent Expressed
10
20
Average Expression
mature-HCs
young-HCs
HC-prog
central-cells
DV-cells
AP-cells
amp-SCs
mantle-cells
Heat Map
Indv. Cell Heatmap
Differential Expression
Pseudotime Analysis
avg.exp.scaled
B
Neuromast Regeneration Trajectory Analysis
Heatmap
Pseudotime Line Plots
Multi-Genes Pseudotime Line Plots
Central Cell Lineage
HC Lineage
Scaled Gene Expression
Pseudotime
UMAP 1
branching point
Neuromast Regeneration scRNA-seq
Group 1 - positive FC
Group 2 - negative FC
Add or remove clusters:
Select statistical test:
wilcox
Input adjusted p-value cutoff:
0.05
Run Differential Expression
Download Results
Gene.name.uniq
p_val
avg_logFC
pct.1
pct.2
p_val_adj
C
gEAR Profile (atoh1a)
Compare two conditions
homeo, Mantle, mantle-cells
Find marker genes

**Figure S2. Overview of the interactive shiny application.**

(A) Homepage to the interactive shiny application for neuromast regeneration analysis. Options for different visualization methods are shown in separate tabs. https://piotrowskilab.shinyapps.io/neuromast_regeneration_scRNA-Seq_pub_2021/

(B) The tab ‘Pseudotime Analysis’ links to interactive shiny app for the neuromast pseudotime analysis with the example of the tab used to graph lineage line plots.

(C) gEAR profile associated with the manuscript (https://umgear.org/p?l=e0d00950) (Orvis et al., 2021). *atoh1a* expression is represented. The portal, in addition to browsing gene expression in the datasets associated with the manuscripts, allows users to explore the data using analysis tools. Here shown is a comparison between HCs and Mantle cells in control condition and an example of marker genes for the different clusters.

### Figure S3

A

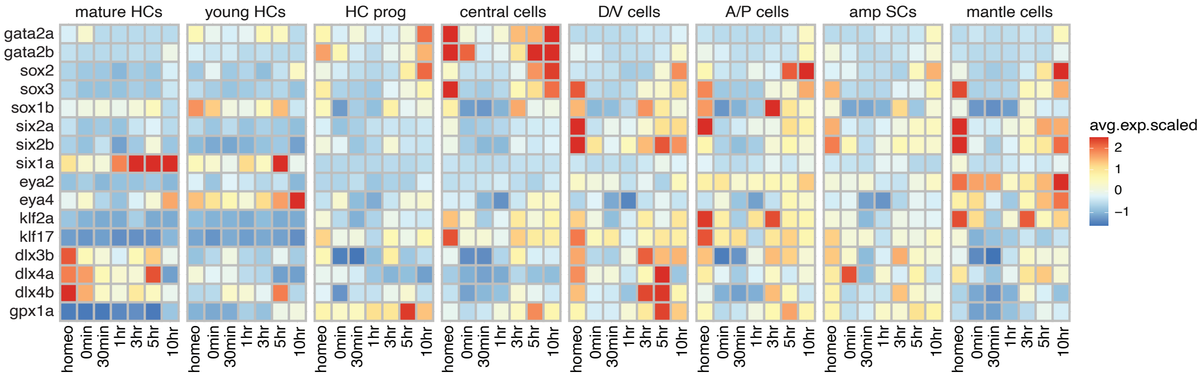
mature HCs
young HCs
HC prog
central cells
D/V cells
A/P cells
amp SCs
mantle cells
gata2a
gata2b
sox2
sox3
sox1b
six2a
six2b
six1a
eya2
eya4
klf2a
klf17
dlx3b
dlx4a
dlx4b
gpx1a
homeo
0min
30min
1hr
3hr
5hr
10hr
avg.exp.scaled
2
1
0
−1

B

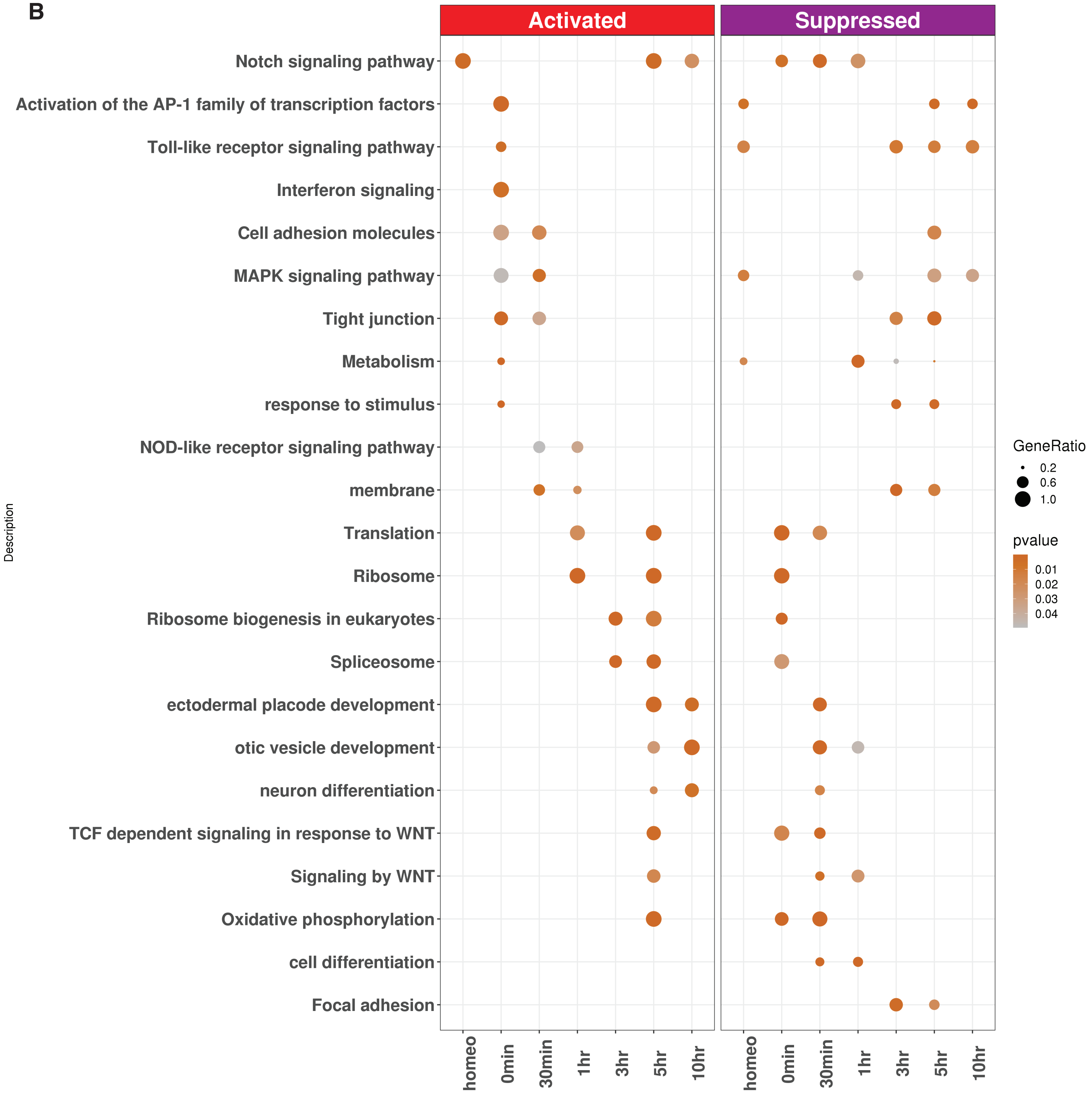
Activated
Suppressed
Description
Notch signaling pathway
Activation of the AP-1 family of transcription factors
Toll-like receptor signaling pathway
Interferon signaling
Cell adhesion molecules
MAPK signaling pathway
Tight junction
Metabolism
response to stimulus
NOD-like receptor signaling pathway
membrane
Translation
Ribosome
Ribosome biogenesis in eukaryotes
Spliceosome
ectodermal placode development
otic vesicle development
neuron differentiation
TCF dependent signaling in response to WNT
Signaling by WNT
Oxidative phosphorylation
cell differentiation
Focal adhesion
homeo
0min
30min
1hr
3hr
5hr
10hr
GeneRatio
0.2
0.6
1.0
pvalue
0.01
0.02
0.03
0.04

**Figure S3. Expression of progenitor/stem cell maintenance/placodal genes and overview of enrichment terms. Related to Figure 2.**

(A) Heatmap of the average expression of downregulated of progenitor/stem cell maintenance/placodal genes across eight cell types during hair cell regeneration.

(B) Overview of gene enrichment terms for up- and downregulated per time point compared to all the other time points.

### Figure S4 0min upregulated

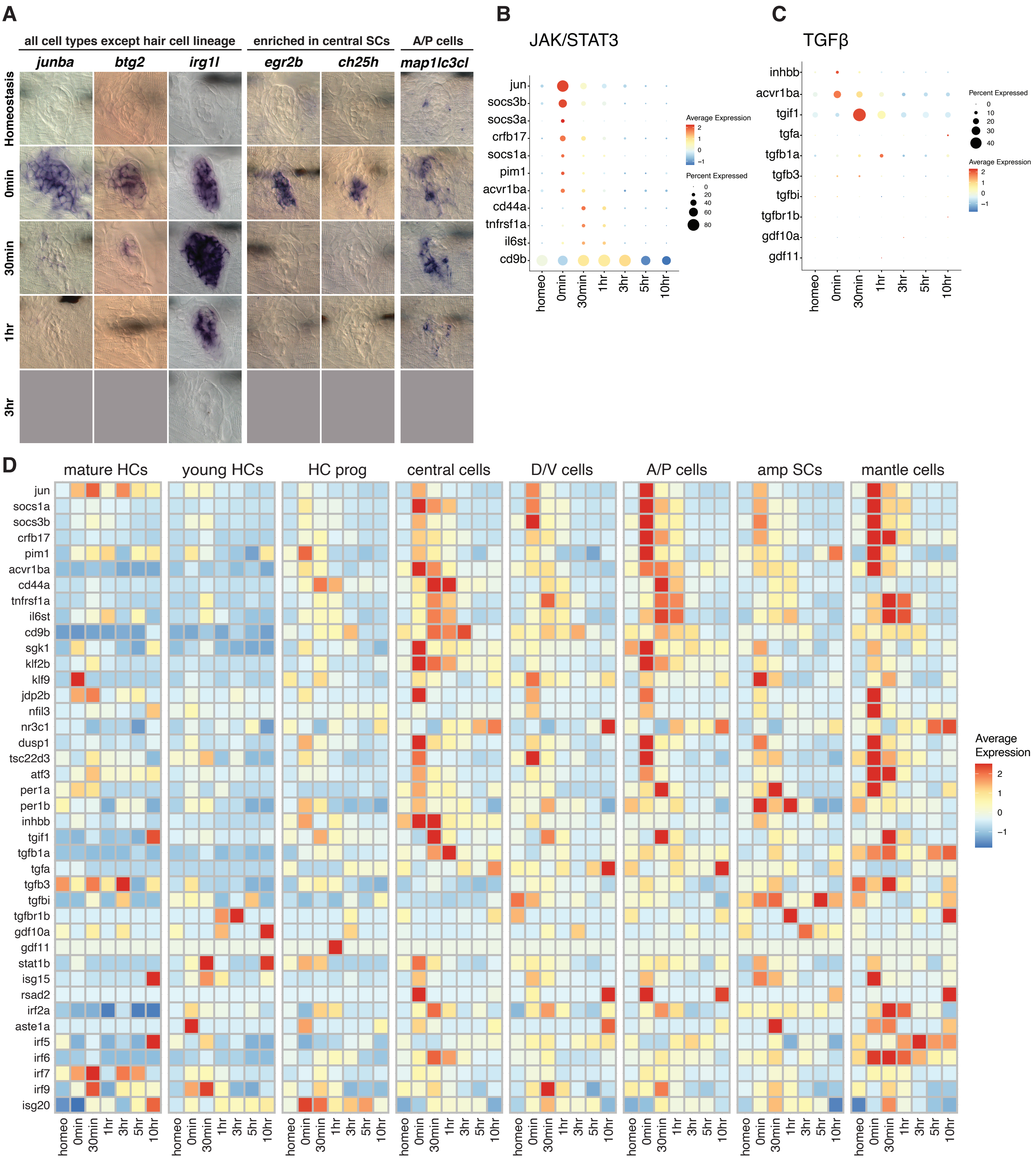

**Figure S4. ISH validation, dot plots and heat maps of genes upregulated at 0min. Related to Figure 3.**

(A) ISH images of genes upregulated at 0min after hair cell death in either all support cells, mostly central support cells or A/P cells.

(B) Dot plot of the average expression of Jak/Stat3 pathway members during hair cell regeneration.

(C) Dot plot of the average expression of TGFβ pathway members during hair cell regeneration.

(D) Heatmap of the average expression of the glucocorticoid, Jak/Stat3 signaling, TGFβ signaling and interferon signaling genes shown in Figure 3N-O during hair cell regeneration.

**Figure S5  heat map of genes in dot plots**

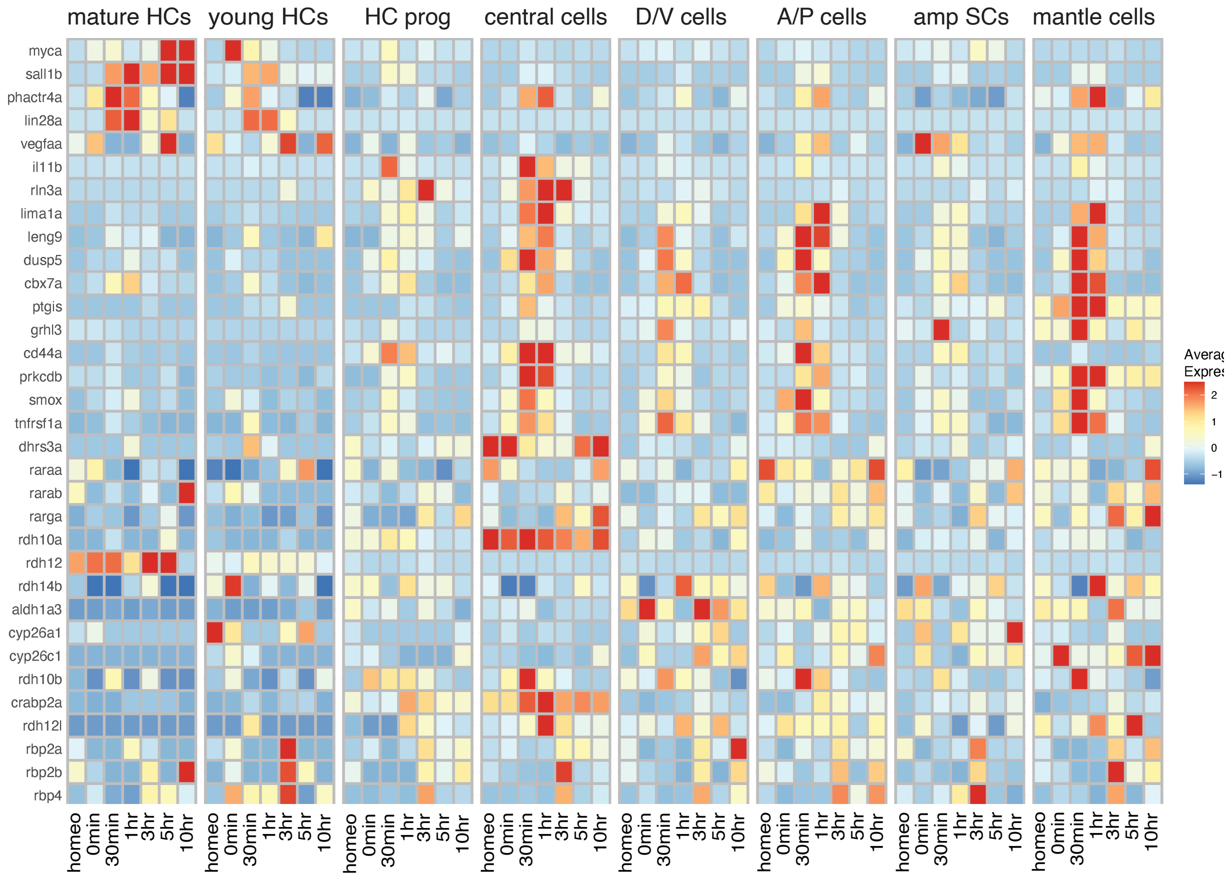

**Figure S5. Related to Figure 4.**

Heatmap of the average expression of the regeneration enriched genes shown in Figure 4H-I and RA signaling genes in Figure 4J-K during HC regeneration.

#### Figure S6

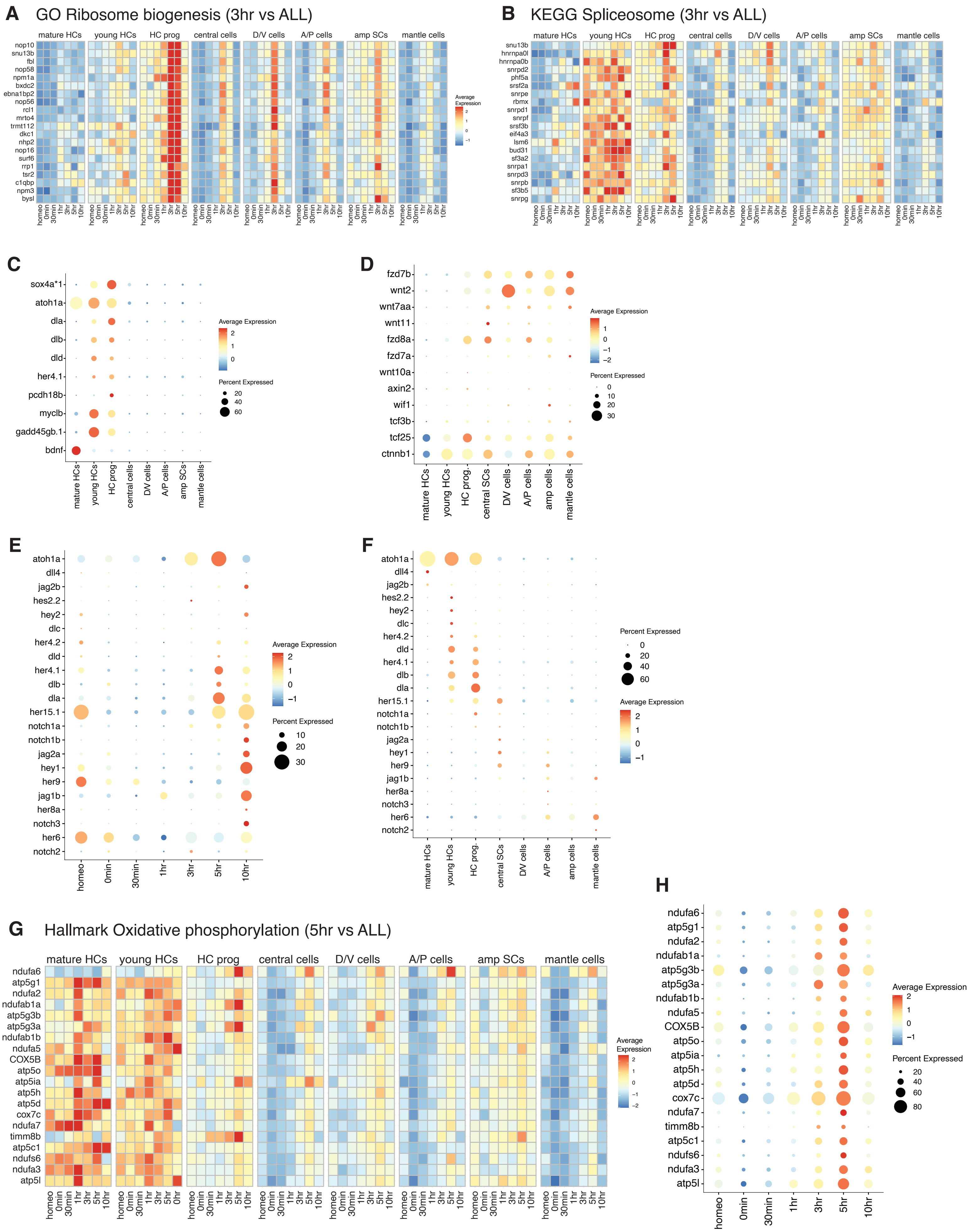
A
GO Ribosome biogenesis (3hr vs ALL)
mature HCs
young HCs
HC prog
central cells
D/V cells
A/P cells
amp SCs
mantle cells
nop10
snu13b
fbl
nop58
npm1a
bxdc2
ebna1bp2
nop56
rcl1
mrto4
trmt112
dkc1
nhp2
nop16
surf6
rrp1
tsr2
c1qbp
npm3
bysl
homeo
0min
30min
1hr
3hr
5hr
10hr
Average Expression
B
KEGG Spliceosome (3hr vs ALL)
snu13b
hnrnpa0l
hnrnpa0b
snrpd2
phf5a
srsf2a
snrpe
rbmx
snrpd1
snrpf
srsf3b
eif4a3
lsm6
bud31
sf3a2
snrpa1
snrpd3
snrpb
sf3b5
snrpg
C
sox4a*1
atoh1a
dla
dlb
dld
her4.1
pcdh18b
myclb
gadd45gb.1
bdnf
Average Expression
Percent Expressed
mature HCs
young HCs
HC prog
central cells
D/V cells
A/P cells
amp SCs
mantle cells
D
fzd7b
wnt2
wnt7aa
wnt11
fzd8a
fzd7a
wnt10a
axin2
wif1
tcf3b
tcf25
ctnnb1
Average Expression
Percent Expressed
mature HCs
young HCs
HC prog.
central SCs
D/V cells
A/P cells
amp cells
mantle cells
E
atoh1a
dll4
jag2b
hes2.2
hey2
dlc
her4.2
dld
her4.1
dlb
dla
her15.1
notch1a
notch1b
jag2a
hey1
her9
jag1b
her8a
notch3
her6
notch2
Average Expression
Percent Expressed
homeo
0min
30min
1hr
3hr
5hr
10hr
F
atoh1a
dll4
jag2b
hes2.2
hey2
dlc
her4.2
dld
her4.1
dlb
dla
her15.1
notch1a
notch1b
jag2a
hey1
her9
jag1b
her8a
notch3
her6
notch2
Percent Expressed
Average Expression
mature HCs
young HCs
HC prog.
central SCs
D/V cells
A/P cells
amp cells
mantle cells
G
Hallmark Oxidative phosphorylation (5hr vs ALL)
mature HCs
young HCs
HC prog
central cells
D/V cells
A/P cells
amp SCs
mantle cells
ndufa6
atp5g1
ndufa2
ndufab1a
atp5g3b
atp5g3a
ndufab1b
ndufa5
COX5B
atp5o
atp5ia
atp5h
atp5d
cox7c
ndufa7
timm8b
atp5c1
ndufs6
ndufa3
atp5l
Average Expression
H
ndufa6
atp5g1
ndufa2
ndufab1a
atp5g3b
atp5g3a
ndufab1b
ndufa5
COX5B
atp5o
atp5ia
atp5h
atp5d
cox7c
ndufa7
timm8b
atp5c1
ndufs6
ndufa3
atp5l
Average Expression
Percent Expressed
homeo
0min
30min
1hr
3hr
5hr
10hr

**Figure S6. Expression of ribosome, spliceosome, hair cell lineage, Wnt-, Notch signaling and OXPHOS genes during HC regeneration. Related to Figure 5.**

(A) Heatmap showing the average expression pattern of ribosome biogenesis-related genes in eight different cell types during hair cell regeneration.

(B) Heatmap of the average expression of spliceosome-term genes (KEGG) in eight different cell types during hair cell regeneration.

(C) Dot plot visualizing the average expression of hair cell lineage genes in eight different cell types (same genes as in Figure 5I).

(D) Dot plot displaying the average expression pattern of Wnt signaling genes in eight different cell types (same genes as in Figure 5K).

(E and F) Dot plots of the average expression pattern of a more comprehensive list of Notch signaling genes during (E) hair cell regeneration and (F) their enrichment in eight different cell types.

(G and H) (G) heatmap and (H) dot plot showing the cell type enriched expression of oxidative phosphorylation-related genes (G) (Hallmark genes) and their upregulation between 3h- 5h after hair cell death (H).

#### Figure S7 heat map of dot plot genes

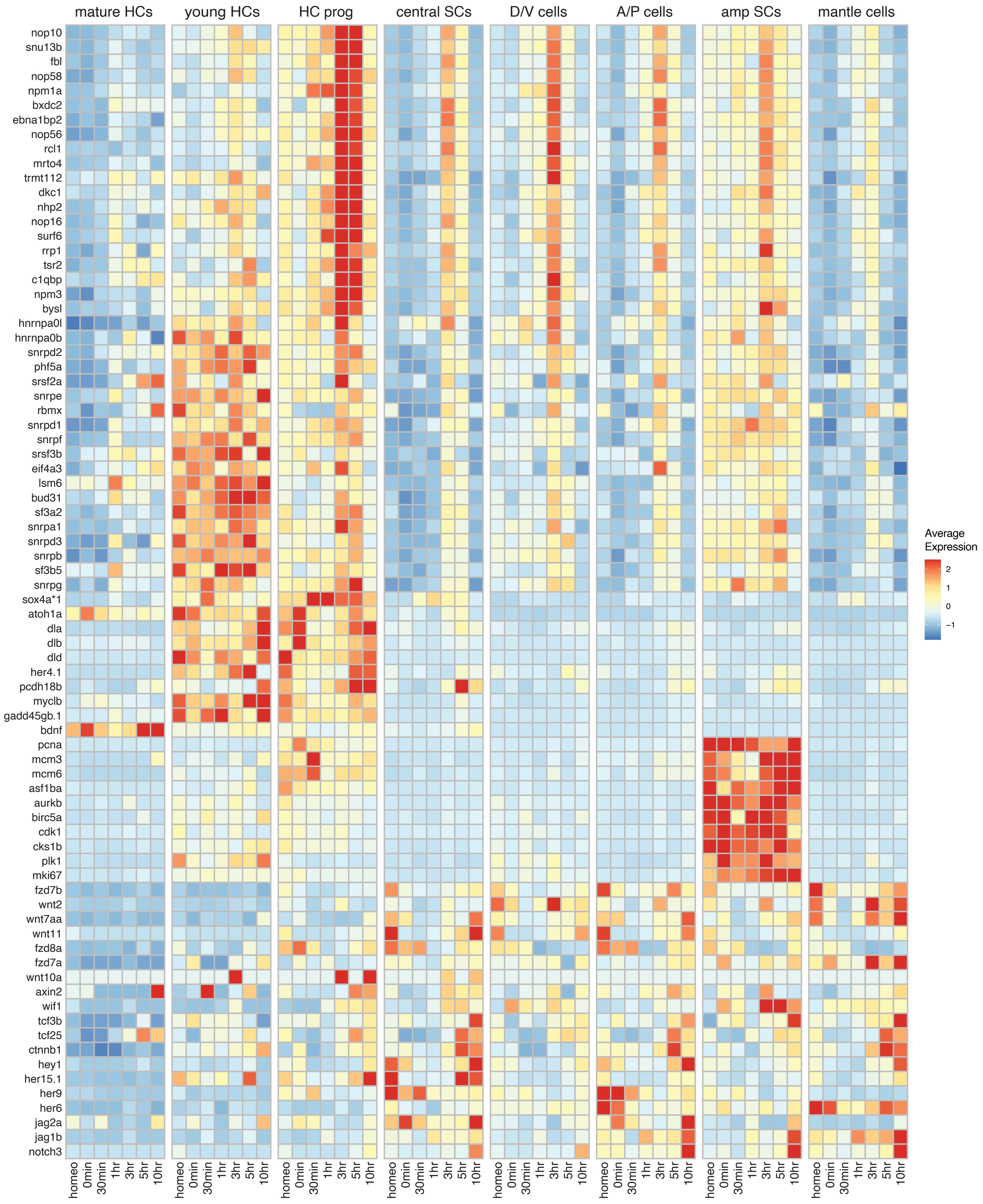

**Figure S7. Related to Figure 5.**

Heatmap visualizing the average expression of ribosome biogenesis, spliceosome, HC lineage, cell cycle, Wnt signaling, and Notch signaling genes shown in the dot plots in Figure 5.

### Figure S8

**A**

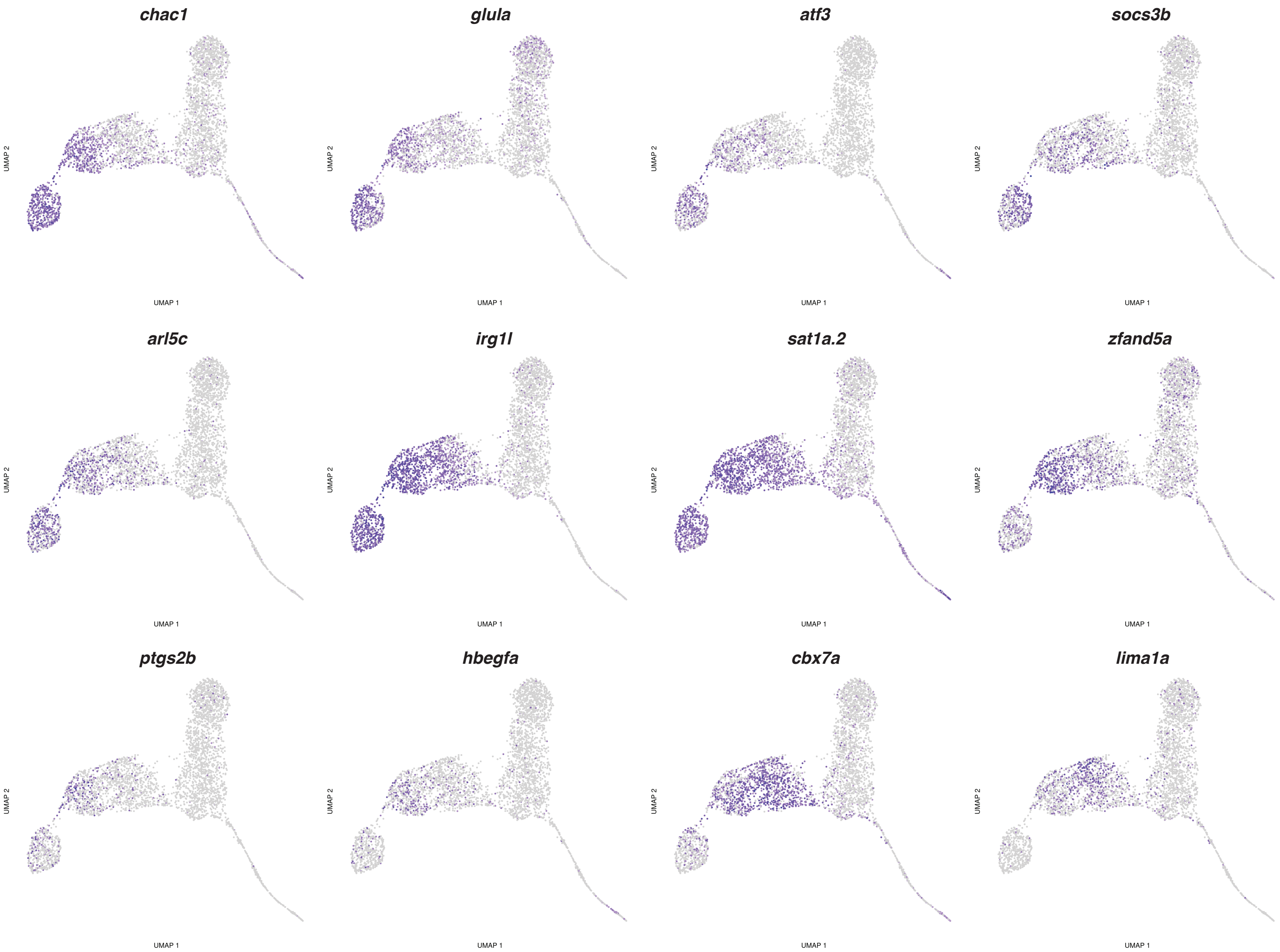

chac1
glula
atf3
socs3b
arl5c
irg1l
sat1a.2
zfand5a
ptgs2b
hbegfa
cbx7a
lima1a
UMAP 1
UMAP 2


**B**

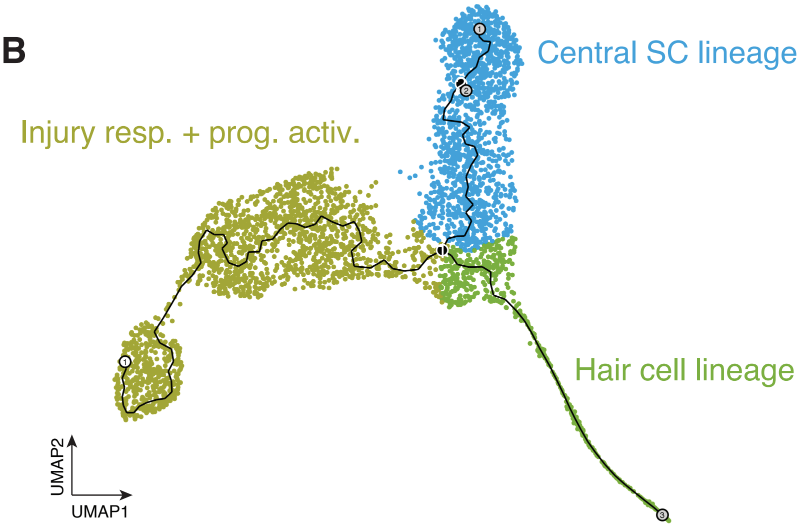

Central SC lineage
Injury resp. + prog. activ.
Hair cell lineage
UMAP2
UMAP1

**Figure S8. Related to Figure 6.**

(A) Gene expression feature plots of representative genes that are enriched in injury response+ progenitor activation clusters (See also in Figure 6H-J).

(B) UMAP showing cell compositions within the Injury response + progenitor activation lineage, central support cell lineage, and hair cell lineage

**Figure S9 Central support cell lineage**

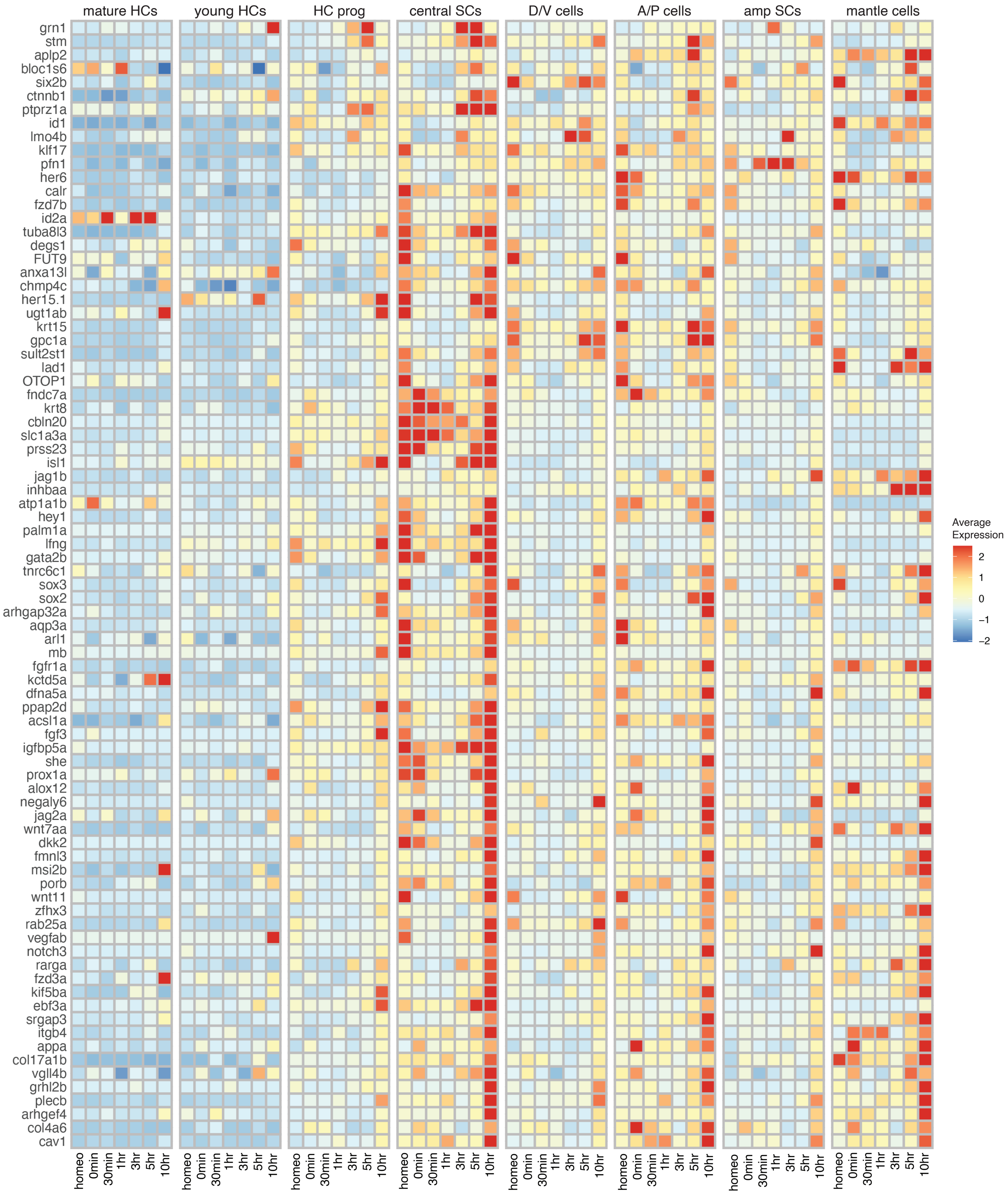

**Figure S9. Central support cell lineage genes. Related to Figure 7.**

Heatmap displaying the average expression of the central support cell lineage genes in clusters 9, 10, and 4 (see also in Figure 7A) in eight different cell types during hair cell regeneration.

### Figure S10  Cell Cycle pseudotime heatmap

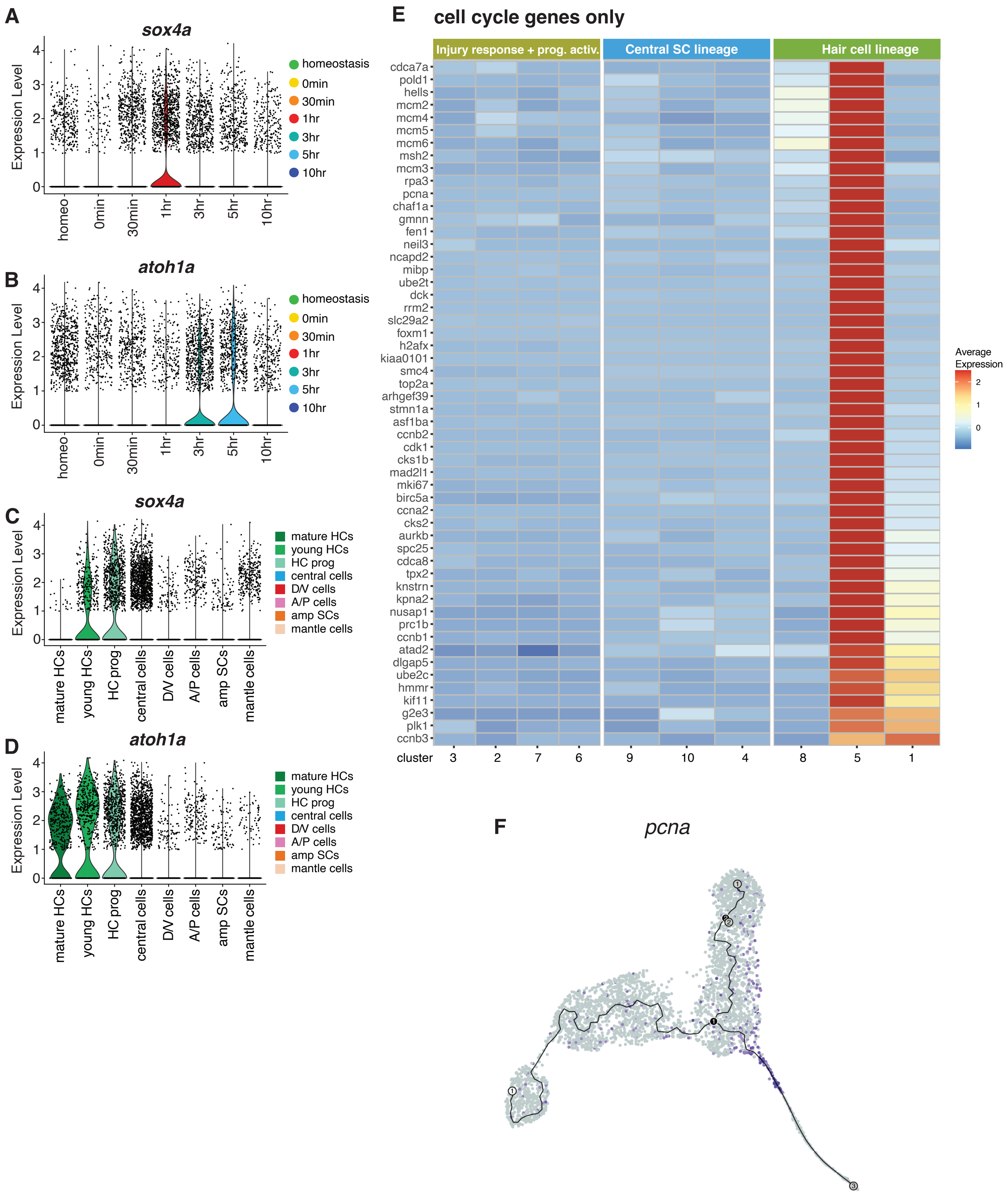

**Figure S10. Related to Figure 8.**

(A-D) Violin plots of *sox4a* (A and C) and *atoh1a* (B and D) across seven different time points of regeneration and eight different neuromast cell types.

(E) Heatmap showing the expression pattern of cell cycle-related genes in the hair cell lineage.

(F) Feature plot of *pcna.*

**Figure S11.**

| | Gene modules | Representative genes | Enrichment terms |
|---|---|---|---|
| | 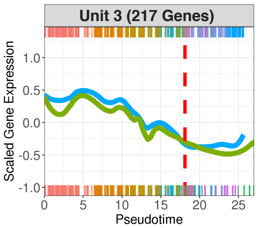 | *nfkbiaa, tnfrsf1a, socs3a, ch25h, cebpb, junba, irg1l, nr4a3, mych, mycb, atf3, socs1a, hbegfa, dusp1, dusp2, gadd45bb, atf4b, jun, nr4a1, fosab* | Response to external biotic stimulus<br>Regulatory region nucleic acid binding<br>Cytokine signaling in immune system<br>Signaling by interleukin<br>MAPK signaling |
| | 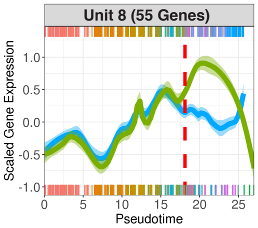 | *RPLs and RPSs* | Translation<br>Metabolism of RNA<br>Ribosome |
| | 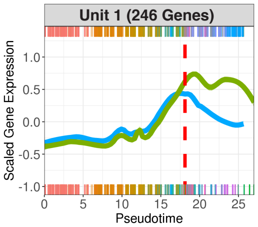 | *nop10, npm1a, nop58,*<br>*nop56, fbl, snu13b*<br>*atp5o, atp5g3b, ndufb8,*<br>*ppa1b, cox7a3*<br>*hnrnpa1b, lsm3, rbm8a*<br>*bud31, lsm8* | Ribosome biogenesis<br>Oxidative phosphorylation<br>Spliceosome |
| | 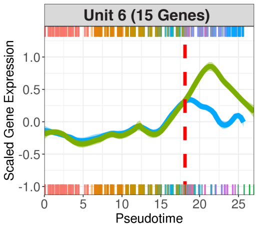 | *mt-co3, mt-co2, mt-cyb,*<br>*mt-atp6, hspd1* | Oxidative phosphorylation<br>Mitochondria part<br>ATP metabolic process |
| Central SC Lineage | 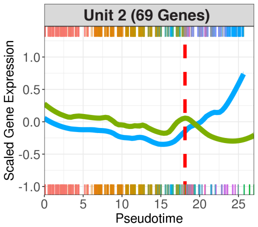 | *sox2, sox3, isl1,*<br>*ebf3a, fndc7a, slc1a3a,*<br>*eya4, hey1, gata2b, rarga,*<br>*stm, id2a, sparc, her15.1,*<br>*cav1* | Sensory organ development<br>inner ear development |
| Hair Cell Lineage | 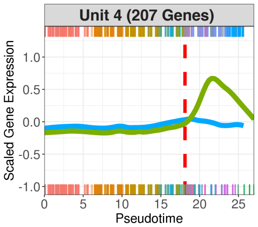 | *rbm24a, myo6b, atoh1b,*<br>*atoh1a, hes2.2, pou4fl*<br>*cdkn2c, mcm3, pttg1, pcna, cdk1*<br>*rfc5, pole2, rnaseh2a, pola2, prim2*<br>*nusap1, h2afva, ccnb1, msh6, suv39h1b*<br>*rfc5, primpol, dek, psmc3ip, msh6* | Cell cycle<br>Cell cycle, Mitotic<br>DNA replication<br>Chromosome organization<br>DNA metabolic process |
| Hair Cell Lineage | 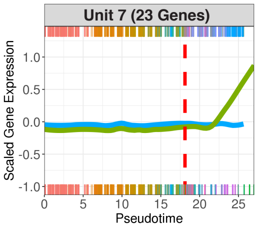 | *efhc2, gpm6ab, nsfa, stmn1b*<br>*snap25a, kif1aa*<br>*emx2, foxj1b*<br>*gapdhs, eno1a, aldocb* | Neuron development<br>Neuron projection development<br>inner ear receptor cell stereocilium organization<br>Glycolysis |
| Hair Cell Lineage | 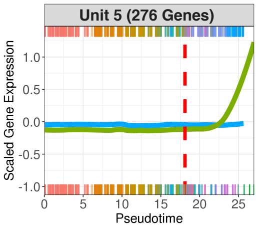 | *tekt3, tekt1, otofa, otofb,*<br>*dnajc5b, s100t, camk2n1a*<br>*cluap1, ttc26, ift57, csnk1e, tctex1d2*<br>*lrrc6, ift88, gas8, dnaaf4, ccdc40*<br>*cfap20, ttc25, lmo7b, catip*<br>*gstp2, sms, gsto2, gstp1, gpx1b* | Cilium Assembly<br>Organelle biogenesis and maintenance<br>epithelial cilium movement<br>Specification of symmetry<br>Glutathione metabolism |

**Figure S11. Gene module (unit) analysis based on the pseudotime analysis. Related to Figure 9.**

Left column: line plots showing the scaled gene expression profiles of genes in each unit ordered along pseudotime (x-axis). Cells belonging to the central support cell lineage are shown as tics on the top x-axis of the graph, whereas cells of the hair cell lineages are represented by tics along the bottom x-axis. Red stitch lines indicate the branch-point between the central support cell and hair cell lineages (Figure 6). Middle column: representative genes in each unit. Right column: representative enrichment terms for each unit.
